# APC couples neuronal mRNAs to multiple kinesins, EB1 and shrinking microtubule ends for bidirectional mRNA motility

**DOI:** 10.1101/2022.07.01.498380

**Authors:** Sebastian J. Baumann, Julia Grawenhoff, Elsa C. Rodrigues, Silvia Speroni, Maria Gili, Artem Komissarov, Sebastian P. Maurer

## Abstract

Understanding where in the cytoplasm mRNAs are translated is increasingly recognised as being as important as knowing the timing and level of protein expression. mRNAs are localised via active motor-driven transport along microtubules (MTs) but the underlying essential factors and dynamic interactions are largely unknown. Using biochemical in vitro reconstitutions with purified mammalian proteins, multi-colour TIRF-microscopy (TIRF-M), and interaction kinetics measurements, we show that adenomatous polyposis coli (APC) enables kinesin-1- and kinesin-2-based mRNA transport, and that APC is an ideal adaptor for long-range mRNA transport as it forms highly stable complexes with 3’UTR fragments of several neuronal mRNAs (APC-RNPs). The kinesin-1 KIF5A binds and transports several neuronal mRNP components such as FMRP, PURα, and mRNA fragments weakly, whereas the transport frequency of the mRNA fragments is significantly increased by APC. APC-RNP-motor complexes can assemble on MTs, generating highly processive mRNA transport events. We further find that EB1 recruits APC-RNPs to dynamically growing MT ends and APC-RNPs track shrinking MTs, producing MT minus-end-directed RNA motility due to the high dwell times of APC on MTs. Our findings establish APC as a versatile mRNA-kinesin adaptor and a key factor for the assembly and bidirectional movement of neuronal transport mRNPs.

## Introduction

The spatial control of gene expression via local translation of mRNAs is an essential prerequisite for the maintenance of cell polarity (Coles and Bradke, 2014; Litman et al., 1993), cell development, cell motility (Kislauskis et al., 1997; Mili et al., 2008) and the co-translational assembly of protein complexes (Moissoglu et al., 2020). In mammalian neurons, hundreds of mRNAs need to be distributed via microtubule-dependent, motor protein-driven transport in the form of small packages (Batish et al., 2012; Park et al., 2014) into a complex network of dendritic branches and the axon until the finest filopodia tips (Buxbaum et al., 2015; Tushev et al., 2018; Wang et al., 2020). As cells control intracellular cargo distribution through a combination of MT-associated proteins (MAPs) and MT post-translational modifications (PTMs) which enable or block the activity of distinct motors (Janke and Magiera, 2020; Kapitein and Hoogenraad, 2015; Monroy et al., 2020), it is expected that the complete localisation process of an mRNA requires either several different adaptors for different motors or a versatile adaptor that can bind several motors (Rodrigues et al., 2021). For instance in *Xenopus laevis*, a combination of a kinesin-1 and a kinesin-2 motor is required to localise Vg1 mRNA (Messitt et al., 2008).

Arriving at a destination such as filopodia (Leung et al., 2018) or axonal branch points (Wong et al., 2017), mRNAs further need to be deposited or “anchored” at cellular structures of the cell cortex at locations where the encoded protein is needed. To date, a major mechanism known to deposit polarity factors at cell cortical elements is microtubule end-binding protein (EB)-dependent MT plus-end-tracking (Bieling et al., 2007; Feierbach et al., 2004; Maurer et al., 2012). MT plus-end-tracking creates transient density waves of plus-end-tracking proteins around polymerising MT ends (Akhmanova and Steinmetz, 2008) and these high local concentrations are thought to drive the formation of polarity factor complexes at the cortex (Martin and Chang, 2003; van Haren et al., 2014). Hence, the question arises whether a mechanism exists that enables mRNA deposition at the cortex, in a similar way as for protein deposition, through MT plus-end–tracking-dependent, concentration-driven complex assembly. While a trailing of mRNPs behind polymerising MT ends has been observed in axonal growth cones (Leung et al., 2018), a mechanism that could enable mRNP MT plus-end-tracking or trailing remains unknown.

Identifying a versatile adaptor which on one hand specifically recognises localisation elements of mRNAs and on the other hand can also bind to different kinesins and eventually EB proteins would close a major gap in our understanding of how mRNPs could be distributed in mammalian cells. While we identified APC as an mRNA adaptor that directly binds G-rich 3’UTR elements and the kinesin-2 KIF3AB-KAP3 (“KIF3ABK”) with high affinities (Baumann et al., 2020), more evidence exists from methods not proving direct interactions which hints towards and alternative, kinesin-1-based transport route for APC (Ruane et al., 2016). This is of high interest, as the kinesin-1 KIF5 was long thought to be a major motor for mRNP transport in different mammalian cell types (Dictenberg et al., 2008; Kanai et al., 2004; Scarborough et al., 2021) but a minimal adaptor sufficient to couple mRNAs to KIF5 is not known. Further, APC interacts with EB proteins (Fig.S1a, Su et al., 1995) and tracks polymerising MT plus ends (Mimori-Kiyosue et al., 2000), which predicts that APC-mRNPs potentially localise to polymerising MT ends. mRNP plus-end-tracking would be an entirely new mechanism to concentrate mRNPs locally, which could promote concentration-driven mRNP remodelling or anchoring of mRNPs at cortical cell structures for subsequent local protein production.

Using a combination of biochemical in vitro reconstitutions with pure proteins and RNA fragments (Grawenhoff et al., 2022), in vitro motility assays, in vitro microtubule dynamics assays (Bieling et al., 2010) and surface plasmon resonance (SPR) interaction measurements, we show for the first time that APC couples neuronal mRNAs to the kinesin-1 KIF5A and the MT plus-end-tracking protein EB1. This results in kinesin-1-driven, fast, and highly processive plus-end-directed APC-RNP motility and EB1-mediated concentration of APC-RNPs at MT plus ends. In addition, we found that lattice-diffusing APC-RNPs can track shrinking MT ends resulting in minus end-directed RNA motility. Thus, APC is a versatile adaptor that links multiple G-rich fragments from at least four different neuronal mRNA-3’UTRs to KIF3ABK, KIF5A and EB1, enabling bidirectional mRNP movements and potentially permitting APC-RNPs to access a wide range of dendritic and axonal segments.

## Results

### KIF5 transports APC-RNPs

Prompted by a published loss-of-function and co-IP study showing that mammalian KIF5A, KIF5B and KIF5C interact with the APC C-terminus (Ruane et al., 2016), we set out to test whether APC directly interacts with KIF5 and whether APC is sufficient to couple neuronal mRNAs to KIF5. As we have previously shown that APC directly links 3’UTR-mRNA fragments to the KIF3ABK trimer (Baumann et al., 2020), this would expand the interaction repertoire of APC, establishing it as a versatile mRNA-motor adaptor. Using biochemical in vitro reconstitutions in combination with TIRF-M (Grawenhoff et al., 2022), we compared the ability of the purified full-length mouse KIF5AA homodimer and the mouse KIF3ABK trimer (KIF5AA, Fig.S1b) to transport reconstituted complexes of APC and β2βtub_wt_ 3’UTR-mRNA fragments (APC-RNPs, Fig.S1c) at experimental conditions in which APC is mostly dimeric and binds one RNA per APC monomer (Baumann et al., 2020). We found that KIF5AA homodimers processively transport APC-RNPs (Fig.1a and movie S1) and that transported complexes readily contain both APC and β2Btub_wt_ RNAs (Fig.1b). At identical RNA:APC ratios, KIF3ABK and KIF5AA transport identical amounts of RNA (Fig.1c), but their motile behaviour differs significantly. As a measure for the motile behaviour of KIF5AA and KIF3ABK-based mRNA transport complexes, we used the confinement ration, which measures how efficient a displacement is in moving away from the initial position (Meijering et al., 2012). The movement of KIF3ABK is less confined than that of KIF5AA (Fig.1a&d), which is reflected in short diffusive periods within tracks of KIF3ABK-mediated RNP transport (Fig.1a). At the same time, KIF5AA-based RNP transport exhibited an almost 2-fold lower dwell time on MTs (Fig.1e), and even though the velocity of KIF5AA-mediated transport is 33% higher (Fig.1f), the total distance travelled is several µm shorter, establishing KIF3-APC complexes as the more processive mRNA transporter (Fig.1g).

**Fig. 1.**
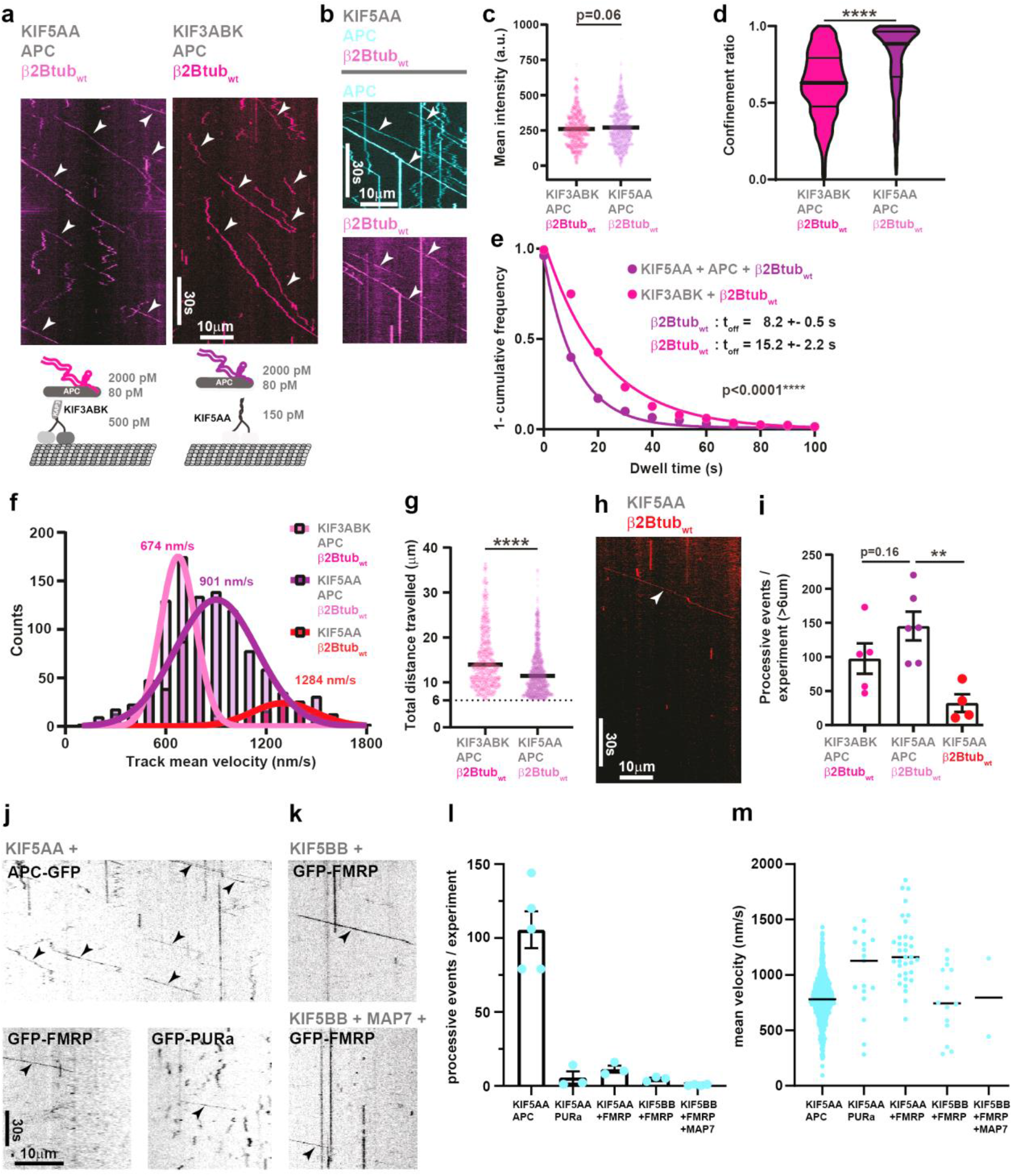
APC links β2βtub-RNA to kinesin-2 and kinesin-1 for processive transport. **(a)** KIF3ABK and KIF5AA transport APC/647-β2Btub_wt_ complexes. Kymographs showing comparable numbers of processive 647-β2Btub_wt_ transport events driven by kinesin-2 (pink) and kinesin-1 (magenta) in the presence of APC. Cartoons depict the components of each minimal transport system and indicate concentrations used for both systems. **(b)** 150 pM KIF5AA transports RNPs containing 80 pM APC-TMR and 2 nM 647-β2Btub_wt_. A kymograph showing multiple co-run events of APC-TMR and 647-β2Btub_wt_ complexes, indicated by white arrowheads. **(c)** Quantification of 647-β2Btub_wt_ fluorescence mean intensities for KIF3ABK+APC+647-β2Btub_wt_ and KIF5AA+APC+647-β2Btub_wt_ transport complexes. **(d)** Violin plot depicting confinement ratio of processive β2Btub_wt_ run events for kinesin-2 (pink)- or kinesin-1 (magenta)-driven transport in the presence of APC. **(e)** Dwell times of 647-β2Btub_wt_ in the presence of KIF3ABK+APC or KIF5AA+APC. **(f)** Histogram showing mean velocities of processive β2Btub_wt_ run events for kinesin-2 (pink)- and kinesin-1 (magenta)-driven transport in the presence of APC as well as for kinesin-1-driven transport in the absence of APC (red). Pink, magenta and red curves represent Gaussian fits for mean velocity distributions **(g)** Scatter plot showing the total distance that 647-β2Btub_wt_ RNAs travel in the presence of KIF3ABK+APC or KIF5AA+APC. **(h)** Kymograph showing the KIF5AA-driven 647-β2Btub_wt_ transport in the absence of APC. The cartoon depicts the components and indicates concentrations used in the experiment. (**i)** Quantification of processive β2Btub_wt_ run events for kinesin-2 (pink)- or kinesin-1 (magenta)-driven transport in the presence of APC as well as for kinesin-1 driven transport in the absence of APC (red). To exclude 647-β2tub_wt_ diffusion events, only processive events > 6 µm are considered. Error bars indicate SEM. **(j)** Kymographs showing KIF5AA (150 pM)-mediated transport of APC-mGFP (80 pM), mGFP-FMRP (800 pM) and mGFP-PURα (800 pM). **(k)** Kymographs showing KIF5BB (150 pM)-mediated transport of mGFP-FMRP (800 pM) in the absence and presence of MAP7 (10000 pM), respectively. **(l)** Quantification of APC-mGFP, mGFP-FMRP and mGFP-PURα transport events at 150 pM KIF5AA and mGFP-FMRP transport at 150 pM KIF5BB in the absence and presence of MAP7, respectively. **(m)** Mean track velocities of APC-mGFP, mGFP-FMRP and mGFP-PURα transport events. Median velocities are indicated. Statistical significance in (h) was evaluated with unpaired, two-tailed t-tests and in (c-f and l) with unpaired Mann-Whitney tests. Horizontal bars in (c, e and l) represent the median. Horizontal bars in (d) represent median and quartiles.

As shown previously (Baumann et al., 2020), KIF3 has essentially no RNA binding activity under our experimental conditions. *Drosophila* KIF5, however, can directly bind U-rich elements (Dimitrova-Paternoga et al., 2021). To test the specificity of KIF5-RNA interactions, we investigated whether mammalian KIF5AA could also transport a β2Btub_wt_ 3’UTR-mRNA fragment containing a high fraction of G residues (Fig.S1c). We found that KIF5AA can transport this RNA as well (Fig.1h and movie S1), even though with a 5-fold reduced efficiency compared to complexes also containing APC (Fig.1i). Comparing the velocities of β2Btub_wt_ RNA transport by KIF3ABK-APC, KIF5AA-APC and KIF5AA (Fig.1f) surprisingly revealed that KIF5AA-based transport in the absence of APC is the fastest RNA transport mode among the three options.

KIF5 motors were shown to interact with several neuronal transport mRNP components such as the RNA-binding proteins (RBPs) FMRP and PURα either directly or indirectly (Kanai et al., 2004; Zhao et al., 2020). Inspired by the diversity of observed KIF5AA interactions, we next tested whether KIF5AA can also directly bind and transport other mRNP components such as the aforementioned canonical RBPs. As challenges with protein biochemistry forced us to work with mGFP-labelled RBPs, we compared their transport efficiency to mGFP-labelled APC. mGFP-FMRP and mGFP-PURα (Fig.S1d) transport by KIF5AA was only detectable at concentrations ∼10-fold above the concentration needed to observe robust APC-mGFP transport (Fig.1j&l and movies S2-3). Given that a recent study observed a significantly higher association of FMRP with KIF5B vs. KIF5A in pull-down assays using murine brain lysate (Zhao et al., 2020), we also purified full-length, unlabelled KIF5BB (Fig.S1d) and tested its ability to transport mGFP-FMRP. We found that besides KIF5AA, also KIF5BB is not able to transport mGFP-FMRP efficiently in vitro (Fig.1k). The addition of active, full-length MAP7 (Fig.S1d-e), a known activator of KIF5 (Ferro et al., 2022; Hooikaas et al., 2019; Monroy et al., 2018), could not increase the observed low FMRP transport efficiency. We hence conclude that the increased ability of KIF5B to pull FMRP from brain lysate (Zhao et al., 2020) requires additional factors which were not present in our in vitro reconstitution assay and are yet to be identified.

Similar to the high velocity of KIF5AA-mediated RNA transport in the absence of APC (Fig.1f), the transport velocity of FMRP was also significantly higher than that of APC (Fig.1m), hinting towards a distinct transport mode of APC and APC-RNPs. These data further show that although KIF5AA can bind several neuronal transport mRNP components, its affinity to APC is likely orders of magnitude higher, positioning APC as the primary factor for neuronal mRNA-motor coupling among the investigated factors.

### APC-RNA complexes are highly stable

While the preceding data establish APC as a versatile, potential neuronal mRNP transport adaptor, it is not known how stable APC-RNA complexes are. Given that RNP complex stability is a crucial requirement for long-range transport processes, we aimed to shed light on APC-RNA interaction dynamics and affinities using SPR interaction kinetics. To this end, we purified an APC fragment containing a Twin-Strep-tag® and the basic MT-binding domain (“APC-basic”, Fig.S2a), and obtained four different minimal 3’UTR fragments comprising G-rich APC motifs (Fig.S1c). Three of these fragments originate from mRNAs previously described as APC targets (β-actin, β2Btub and Map1b) (Preitner et al., 2014), whereas one fragment originates from the Camk2α mRNA 3’UTR which was not detected as an APC target previously but contains a G-rich structure (G-quadruplex) recognised by FMRP (Darnell et al., 2001; Goering et al., 2020). We first set out to measure the interaction kinetics of APC and β2Btub_wtmin_ RNA vs. APC and a control RNA (β2Btub_mutmin_ RNA), in which G residues were replaced with other nucleotides (Fig.S1c and Fig.2a&b). While APC shows a moderate association rate to β2Btub_wtmin_ RNA (Fig.2a), its dissociation rates using both a one-state and a two-state interaction model, are in the range of 10^−2^ to 10^−4^/s (Fig.2a & Fig.S2b). APC-β2Btub_wtmin_ RNA complexes are hence stable for minutes, and as such are suitable for longer transport processes needed to deliver mRNP packages to different locations within hundreds of micrometres long neurites. Confirming published TIRF-M and microscale thermophoresis (MST) data (Baumann et al., 2020), there is essentially no interaction detectable between APC and the β2Btub_mutmin_ RNA

**Fig. 2.**
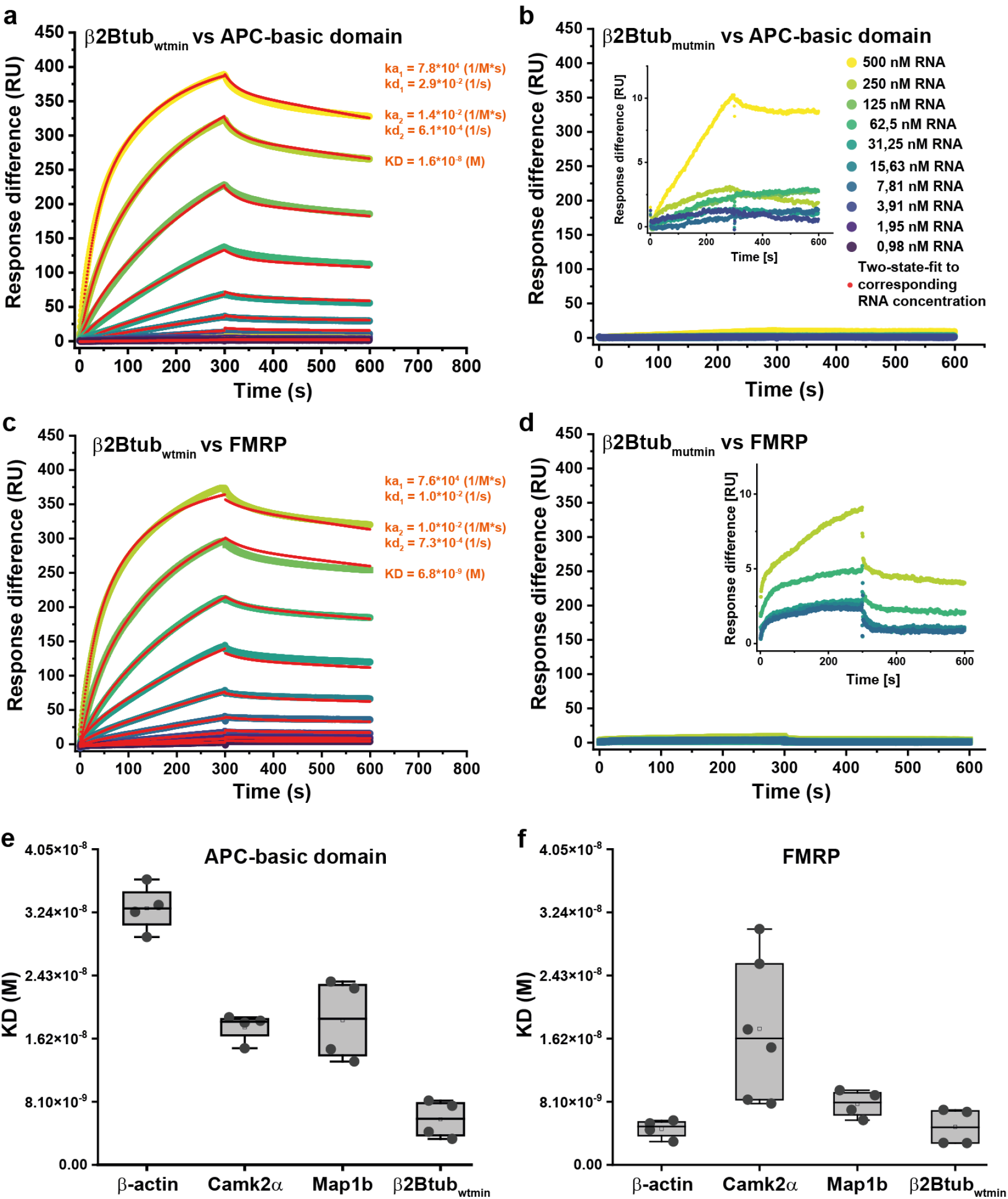
APC-basic domain and FMRP bind G-rich target RNAs with high affinity. **(a & c)** Graphs generated from Biacore experiments showing the binding kinetics (association and dissociation rates) of different concentrations of β2Btub_wtmin_ RNA and APC-basic or FMRP, respectively. Experimentally determined curves were fitted with a two-state binding model. **(b & d)** Graphs generated from Biacore experiments showing the association and dissociation of different concentrations of β2Btub_mutmin_ RNA and APC-basic domain as well as FMRP, respectively. **(e & f)** Plot of APC and FMRP target RNA dissociation constants. Experiments were performed at least in duplicates with two technical replicates each.

FMRP binds G-rich sequences very similar to those recognised by APC and their mRNA target pools at least partially overlap, while the two proteins differ in their functions (Darnell et al., 2011; Dictenberg et al., 2008; Preitner et al., 2014; Reilein and Nelson, 2005). It is hence interesting to understand if both RBPs would compete for the same mRNAs, either for translation regulation or for the assembly of different mRNA transport complexes. Testing β2Btub_wtmin_ and β2Btub_mutmin_ RNAs binding to immobilised FMRP (Fig.S2a) revealed an even higher affinity and similar low off rates of FMRP to β2Btub_wtmin_ RNA (Fig.2c), in comparison to APC, while very little FMRP binding activity was detected for β2Btub_mutmin_ RNA (Fig.2d), similar to APC.

We finally extended the SPR analysis to all four 3’UTR fragments. We found that both APC and FMRP bind all wild type RNAs with nM-range affinities (Fig.2a,c,e-f & Fig.S2b-d). The only difference was detectable for β-actin RNA (Fig.S2d): here, Biacore analysis was only possible with a narrower concentration range, reducing the confidence on the measured difference. We therefore conclude that FMRP and APC would likely compete for the same mRNA localisation motifs but note that seemingly small differences detected here could have a strong effect on mRNA transport. APC, for instance, binds β-actin RNA with 3-fold lower affinity than β2Btub_wt_ RNA. Still, we could show previously that this small difference is sufficient to almost entirely block β-actin mRNA transport in the presence of equimolar amounts of β2Btub_wt_ RNA (Baumann et al., 2020).

### KIF3 and KIF5 do not act synergistically during APC-RNP transport

Having established APC as both a versatile but also efficient mRNA-motor adaptor, we next asked whether APC could employ KIF3 and KIF5 simultaneously, a mechanism which could be beneficial, e.g., for super-processive long-range transport of mRNA along the neuronal axon. Given that KIF3 binds the N-terminal ARM domain of APC via KAP3 (Jimbo et al., 2002), while KIF5 was reported to bind the C-terminus of APC (Ruane et al., 2016) (Fig.S1a), a co-transport scenario would be possible. We hence analysed the motility of complexes containing APC-647 (Fig.S3a) and RNA in the presence of either KIF3ABK (scenario 1), KIF5AA (scenario 3), or both kinesins (scenario 2) (Fig.3a). Qualitatively, the addition of KIF5AA to KIF3ABK-APC-RNA complexes caused little difference in motile behaviour (Fig.3b, (scenario 2)) compared to KIF3ABK-APC-RNA motility (Fig.3b, (scenario1)). The number of processive APC transport events did not increase significantly upon KIF5AA addition (Fig.3c), indicating that the APC concentration was limiting in our experiments. Interestingly, while the velocity distribution of the two-motor scenario (Fig.3d, (scenario 2)) was bimodal with peaks matching the velocities of KIF5AA- and KIF3ABK-based APC-RNA transport, addition of KIF5AA produced a confinement ratio distribution (Fig.3e, (scenario 2)) very similar to that of the pure KIF5AA condition (Fig.3e, (scenario 3). The dwell times (Fig.3f) and the total distance travelled (Fig.3g) of the two-motor scenario (2) are intermediates between the pure KIF3ABK-APC-RNA scenario (1) and the pure KIF5AA-APC-RNA scenario (3). These experiments show that both kinesins are not likely engaged in simultaneous transport of APC, since the association of a single cargo with several motors is expected to decrease its detachment from the MT and thus increase its overall processivity. These results prompted us to reinvestigate which APC domains are bound by the individual kinesin motors.

**Fig. 3.**
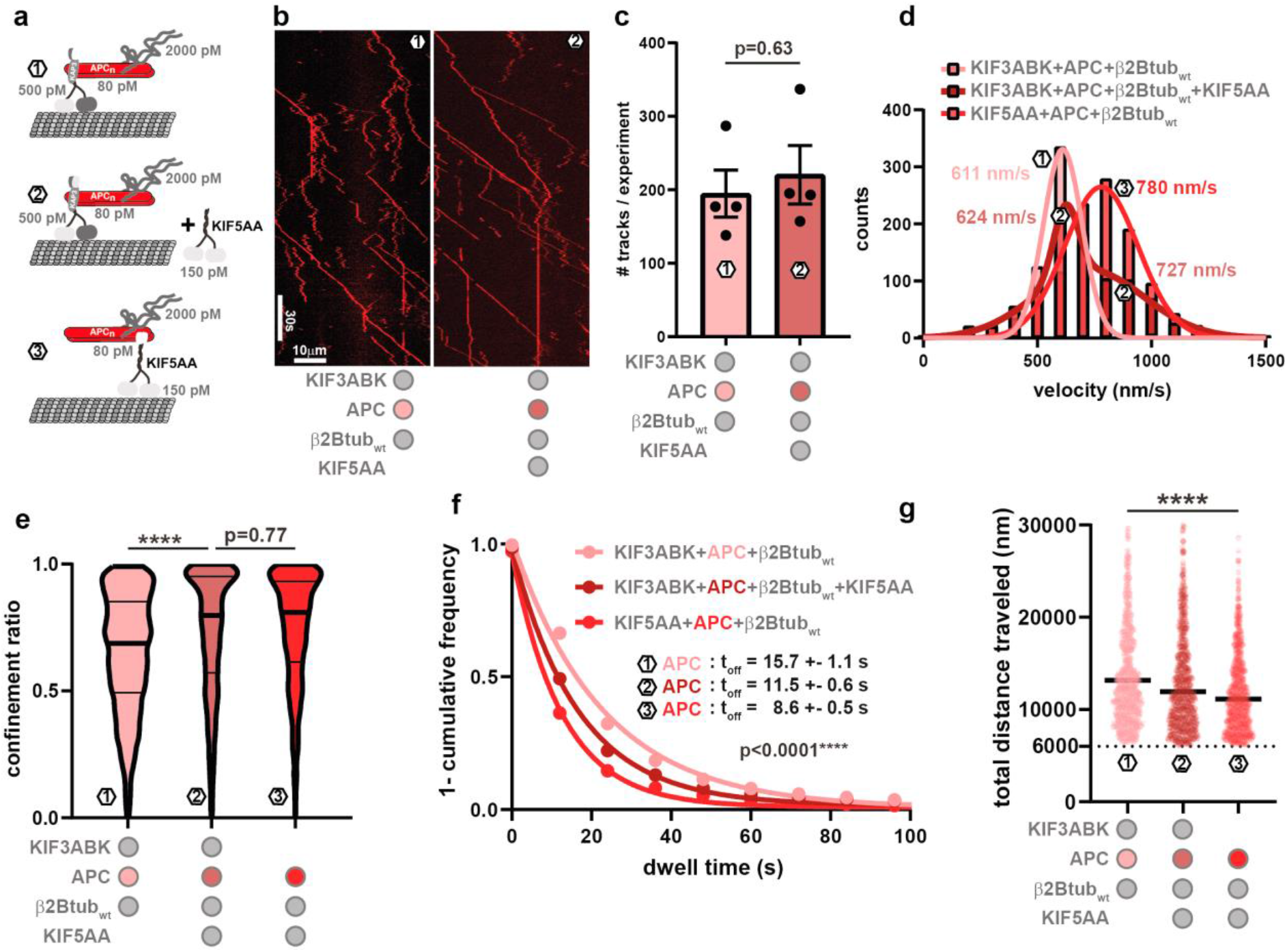
Combined effect of KIF5AA and KIF3ABK on RNA transport. **(a)** Cartoon depicting the experimental setup used to investigate the effect of kinesin motors on APC in different scenarios: (1) KIF3ABK+APC-647; (2) KIF3ABK+APC-647+KIF5AA; (3) APC-647+KIF5AA. Picomolar concentrations of components are indicated. **(b)** Kymographs showing APC-647 in the presence of β2Btub_wt_ and KIF3ABK (left panel), or in the presence of β2Btub_wt,_ KIF3ABK and KIF5AA (right panel). **(c)** Quantification of processive RNP transport events containing APC-647 in the presence of β2Btub_wt_ and KIF3ABK or in the presence of β2Btub_wt,_ KIF3ABK and KIF5AA, respectively. Statistical significance was evaluated with an unpaired, two-tailed t-test. **(d)** Velocity distributions of APC-647 in the presence of KIF3ABK (1), KIF3ABK and KIF5AA (2), or KIF5AA (3). Velocity distribution of (1) and (3) were fitted with a Gaussian curve whereas (2) was fitted with the sum of two Gaussians. **(e)** Violin plot depicting the confinement ratio of transported APC-647 in conditions (1), (2) and (3). Statistical significance was evaluated with an ordinary one-way ANOVA test. **(f)** Dwell times of APC-647 in all three conditions. (e, f) Statistical significance was evaluated with a Kruskal-Wallis test. **(g)** Total distance travelled of APC-647 in all three conditions. Statistical significance was evaluated with an unpaired Mann-Whitney test.

### Both kinesins bind and transport the APC-ARM domain

While the binding of the KIF3ABK trimer or KAP3 alone to the APC-ARM domain was shown with methods reporting direct interactions (Baumann et al., 2020), binding of KIF5 to the APC C-terminus was only tested with methods that cannot discriminate between direct or indirect interactions (Ruane et al., 2016). We hence purified an Alexa647-labelled C-terminal APC fragment (“APC-C”, Fig.S4a), containing the MT-binding region and the proposed KIF5 binding site, but lacking the APC-ARM domain. Comparing the motility of full-length APC with APC-C in the presence of unlabelled KIF5AA showed that the deletion of the ARM domain reduces processive APC transport events, while the number of diffusive events is increased (Fig.4a-c). Almost no KIF3ABK-mediated transport of APC-C was detected, underlining that this motor specifically recognises the ARM region in the N-terminal domain of APC (Fig.4a-c). Of note, the velocity of KIF5AA-mediated APC-C transport is higher than the velocity of full-length APC transport (Fig.4d). This observation led us to speculate that full-length APC might have multiple MT binding sites, which could lead to a transport mode in which APC continues to interact with the MT during transport. This could potentially reduce APC transport velocity but could also increase processivity.

**Fig. 4.**
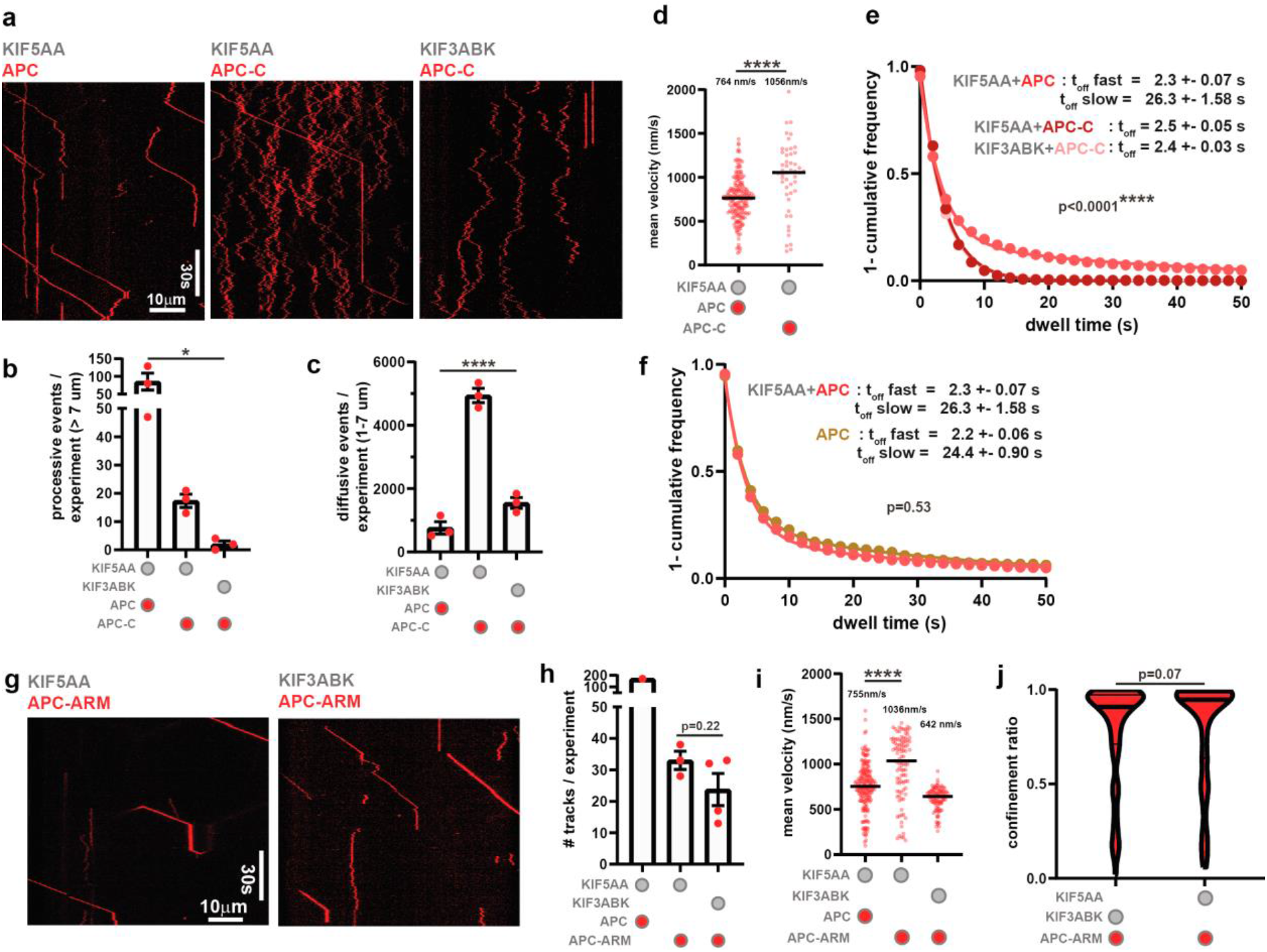
KIF5AA interacts with the APC N- and C-termini. **(a)** Kymographs showing motion behaviour of full-length APC-647 (25 pM) and APC-C-647 (25 pM) in the presence of KIF5AA (150 pM) or KIF3ABK (500 pM). **(b & c)** Quantification of processive (b) and diffusive events (c). Statistical significance was evaluated with one-way ANOVA tests. **(d)** Track mean velocities of full-length APC-647 and APC-C-647 transported by KIF5AA. Statistical significance was evaluated with an unpaired, two-tailed t-test. **(e)** 1-cumulative frequency plot for the determination of diffusive full-length APC-647 and APC-C-647 in the presence of KIF5AA or KIF3ABK. Only events with a track displacement > 1 µm and a maximum distance < 7 µm were considered. Experimental data of APC-C-647 were fitted with a one-phase decay curve, whereas full-length APC-647 was fitted with a two-phase decay curve. Statistical significance between APC-647 and APC-C-647 in the presence of KIF5AA was evaluated with a Mann-Whitney test. **(f)** 1-cumulative frequency plot comparing the dwell times of diffusive APC in the presence or absence of KIF5AA. Experimental data were fitted with two-phase decay curves. Statistical significance was evaluated with a Mann-Whitney test. **(g)** Kymographs showing motion behaviour of APC-ARM-647 (100 pM) in the presence of KIF5AA (150 pM) or KIF3ABK (500 pM). **(h)** Quantification of processive events. Full-length APC-647 (100 pM) transported by KIF5AA serves as control similar to (b). Statistical significance was evaluated with an unpaired, two-tailed t-test. **(i)** Track mean velocities of full-length APC-647 and APC-ARM-647 transported by KIF5AA as well as APC-ARM-647 transported by KIF3ABK. Statistical significance was evaluated with an unpaired, two-tailed t-test. **(j)** Confinement ratio of APC-ARM-647 in the presence of KIF5AA and KIF3ABK. Statistical significance was evaluated with a Mann-Whitney test.

As we would expect that a multivalent attachment of full-length APC to the MT lattice would increase its dwell time, we first tested whether the dwell times of full-length APC and APC-C differ. Since the number of processive events for APC-C is too low for further analysis, we only took diffusive full-length APC and APC-C events into account. To this end, we applied a tracking filter, allowing us to restrict the analysis to APC and APC-C tracks with a displacement between 1 and 7 µm, a distance characteristic for diffusing APC (Baumann et al., 2020). Full-length APC shows a biphasic dwell time with either very short diffusion events of 2.3 s or approximately 10 times longer diffusion events of 26.3 s, indicating that it harbours at least two MT binding sites (Fig.4e). In contrast to full-length APC, APC-C dwell time curves are best fitted with a mono-exponential curve, exclusively showing higher turnover events of 2.5 s, which most likely result from a single MT-binding site. Considering that the dwell times of an “APC only” condition are almost identical to those of the “KIF5AA+APC” condition (Fig.4f), we can rule out convolutions caused by the presence of KIF5AA.

As our data indicates a competition of both kinesin motors for the APC cargo (Fig.3), we next asked whether in addition to KAP3, KIF5AA might also be able to bind the APC-ARM fragment. We found that although KIF5AA transports full-length APC with a higher efficiency, both KIF5AA and KIF3ABK can processively transport the APC-ARM fragment (Fig.4g-h). Intriguingly, as for APC-C, the velocity of KIF5AA-mediated APC-ARM transport was again higher compared to that of full-length APC (Fig.4i). Also, the previously noted diffusive component in KIF3ABK-mediated transport of full-length APC (Fig.1d) was largely removed when APC-ARM was transported (Fig.4j). These data show that both kinesins are likely to compete for the APC cargo to some extent due to their mutual binding to the APC-ARM domain. Further, full-length APC likely binds MTs with an N- and a C-terminal MT-binding site which together affect APC transport by KIF3ABK and KIF5AA: upon removal of either APC terminus, the KIF5AA-based transport velocity of APC increases.

### Transport APC-RNPs can assemble on the microtubule

Neuronal mRNPs exhibit a random Lévy walk; while being non-motile most of the time, mRNPs can diffuse and exhibit bidirectional transport (Song et al., 2018). Such a movement pattern can improve target site finding of randomly located objects (Viswanathan et al., 1999). A mechanism in which kinesin motors push or drag MT lattice-diffusing APC-RNPs would thus offer an attractive new framework for interpreting neuronal mRNP movements. We investigated the movements of Alexa-647-labelled KIF5AA and TMR-labelled APC in dual colour TIRF-M experiments and found examples in which KIF5AA and APC landed simultaneously on MTs, indicating transport RNP assembly before MT binding (Fig.5a, magnified frame on the middle right). Strikingly, we also observed that faster moving KIF5AA encounters lattice-diffusing APC (Fig.5a and movie S4). In some cases, the motors “pick up” lattice-diffusing APC for further transport (Fig.5a, magnified frames on the right, top and bottom), which creates RNP trajectories with stationary rest phases and processive run events, as observed in mammalian neurons (Song et al., 2018; Yoon et al., 2016). Of note, once KIF5AA is loaded with its APC cargo, a decrease in KIF5AA velocity can be observed. Labelling of both KIF5AA and APC enabled us to calculate the ratio of processive APC transport events to processive KIF5AA events which was ∼1:4 (Fig.5b). Considering the high APC labelling ratio of approximately 80% and its propensity to form a dimer (Baumann et al., 2020), we assume that at least one of the APC molecules in each dimer is labelled. Hence, the majority of KIF5AA run events can be considered “cargo-free” run events. We analysed the motility parameters of APC and KIF5AA populations separately. Despite the expected ∼25% overlap of both data sets, we found that the KIF5AA data showed a higher velocity, resembling the velocity of KIF5AA loaded with either RNA, FMRP or APC fragment cargoes (Figs.1&3), while the APC fraction was moving at the characteristic KIF5AA-APC complex velocity (Fig.5c). At the same time, the processive APC data exhibited a 2.5-fold higher dwell time (Fig.5d) than the cargo-free KIF5AA data, and KIF5AA-transported APC travelled several µm farther than KIF5AA alone (Fig.5e), although with a greater diffusive component than that of KIF5AA movements (Fig.5f). In principle, motor oligomerisation by the dimeric APC could cause such changes in motile behaviour. However, we found no correlation between measured KIF5AA signal intensity and velocity (Fig.5g). Thus, motor oligomerisation by APC is unlikely the underlying reason for APC-induced motor slow down.

**Fig. 5.**
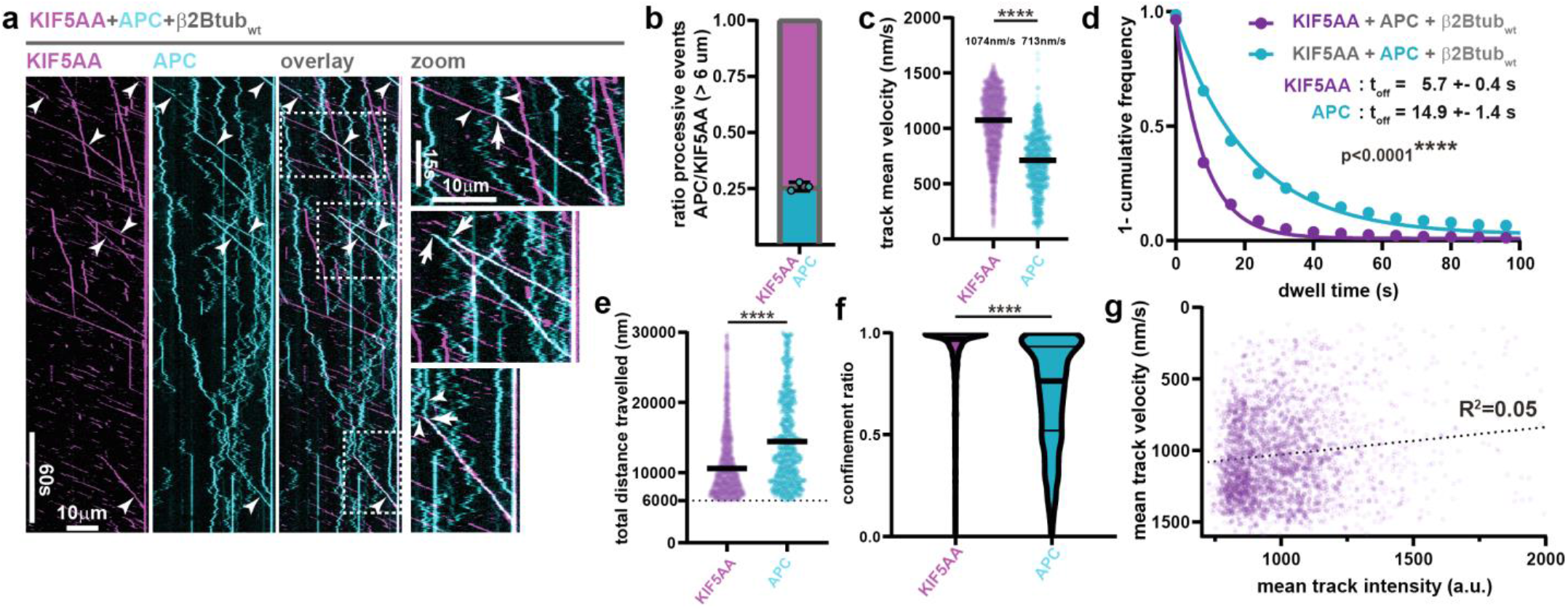
APC changes KIF5AA motility parameters. **(a)** Kymographs showing the motion behaviour of KIF5AA-647 (80 pM) in magenta and APC-TMR (50 pM) in cyan. APC-TMR transported by KIF5AA appears white in the overlay of kymographs. White dotted rectangles indicate zoomed parts showing diffusive APC-TMR being picked up by KIF5AA-647 resulting in processive APC-TMR transport (top and bottom panel) as well as co-landing events of APC-TMR and KIF5AA-647 also resulting in processive APC-TMR transport (middle panel). **(b)** Ratio of processive KIF5AA-647 transporting APC-TMR. Only events > 6 µm were considered. **(c)** Track mean velocities of KIF5AA-647 and APC-TMR populations. Note that about 25% of processive KIF5AA-647 transports APC-TMR. Median velocities are indicated and statistical significance was evaluated with a Mann-Whitney test. **(d)** 1-cumulative frequency plot comparing dwell times of KIF5AA-647 and APC-TMR. Statistical significance was evaluated with a Mann-Whitney test. **(e)** Scatter plot depicting the total distance travelled of APC-TMR and KIF5AA-647. Statistical significance was evaluated with a Mann-Whitney test. **(f)** Violin plot depicting confinement ratios of KIF5AA-647 and APC-TMR. Statistical significance was evaluated with a Mann-Whitney test. **(g)** Scatter plot testing the correlation of KIF5AA-647 mean track velocity and mean track intensity. The dotted line represents simple linear regression. R^2^ indicates R squared.

In summary, this data supports a mechanism in which the long dwell time of APC on the microtubule promotes encounters with KIF5AA which on its own binds only briefly, exhibiting short directional runs. Once KIF5AA-APC complexes assemble on the microtubule, KIF5AA is slowed down, but becomes more processive. As these changes in motile behaviour are not correlated with oligomerisation of the motor protein, and do not occur when any other cargo tested here is transported, we conclude that the simultaneous interaction of APC dimers with the MT and motor likely causes this effect on motor activity.

### APC-RNPs track growing and shrinking microtubule ends

APC is an EB-dependent microtubule plus-end-tracking protein (Mimori-Kiyosue et al., 2000) and APC-dependent mRNA localisation to protrusions of axonal growth cones and NIH/3T3 cells were shown to be of crucial importance for polarisation and migration (Mili et al., 2008; Preitner et al., 2014). We hence asked whether APC could also recruit mRNAs to polymerising microtubule plus ends in addition to being an mRNA-motor adaptor. We purified EB1 and EB1-mGFP (Fig.S4a), and initially used an in vitro TIRF-M assay with dynamically growing microtubules to assess the dynamic behaviour of EB1 and APC. We observed robust, EB1-dependent APC tracking of growing microtubule ends (Fig.6a and movie S5). Unexpectedly, APC could also track shrinking MT ends. As EB proteins recognize a GTP hydrolysis dependent tubulin structure that only occurs during MT assembly, they cannot track depolymerising microtubule ends (Bieling et al., 2007; Maurer et al., 2011, 2012). Hence APC tracks shrinking MT ends independently of EB1 (Fig.6a). The high-affinity interaction between EB1 and APC in solution could further be confirmed with Biacore measurements, showing multi-site binding on both the full APC–C-terminus (“APC-C”, residues 2163-2845) and the APC-basic domain alone (APC residues 2163-2670, Fig.S4a-d). This confirms that the C-terminal half of APC contains multiple EB-binding sites as predicted by the presence of six extended EB-binding SXIP motifs (Jiang et al., 2012) in this region. We employed automated tracking of EB1-mGFP to determine MT growth speed and time of extension periods (Fig.S4e and movie S6). Under our experimental condition, the presence of APC had no effect on MT growth speed or the extension period of a polymerising microtubule (Fig.6b). Addition of β2Btub_wt_ RNA resulted in plus-end-tracking of APC-RNPs (Fig.6c and movies S7&S8), with APC-RNPs also being able to track EB1-free, shrinking microtubules (Fig.6c). Of note, the addition of RNA had no effect on MT growth speed or extension period (Fig.6d). Control experiments show that APC-RNA complexes cannot track MT plus ends in the absence of EB1 (movies S9) and that RNA is recruited to MT plus ends in an APC-dependent manner (movie S10). Given the high-affinity interaction between EB1 and APC, we asked next whether EB1 can affect the dwell time of APC on MTs. We found that both the fast and the slow component of the APC dwell time on paclitaxel stabilised MTs increase with EB1 concentration to values of > 60 s (Fig.6f). The long APC dwell times, even in the absence of EB1, are sufficient to enable processive shrinking-end-tracking of APC-RNA complexes, as APC dwell times can well extend beyond the duration of an MT shrinking event. Upon the addition of KIF3ABK, we observe bidirectional APC-RNA complex movements along MTs (Fig.6g-h), caused by kinesin-dependent, plus-end directed transport and the shrinking-end-tracking behaviour of APC, potentially enabling mRNPs to enter and scan through fine cell protrusions and facilitate mRNA-target site finding by repeated and alternating cycles of MT plus- and minus-end-tracking with active transport (movie S11).

**Fig. 6.**
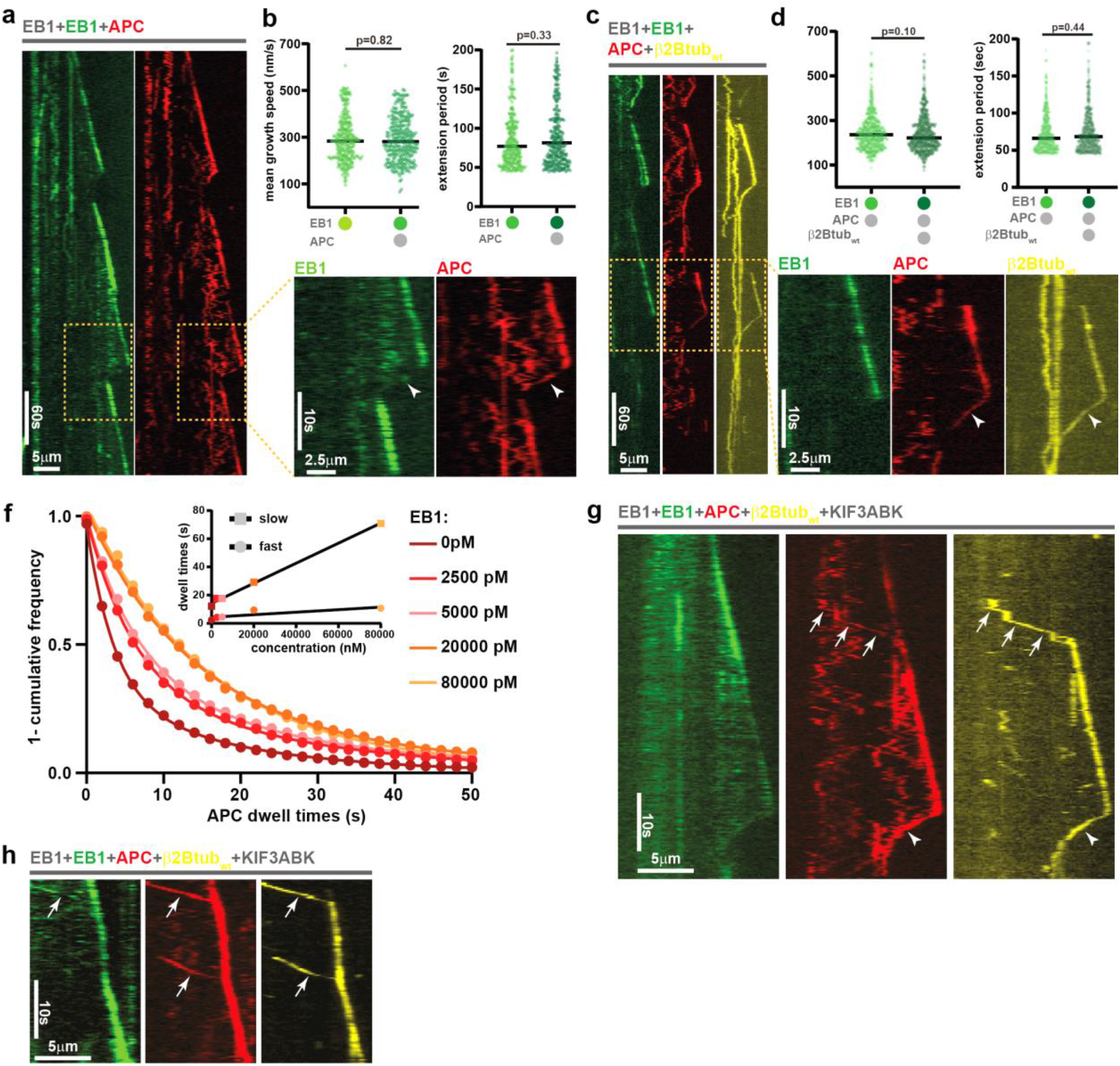
Microtubule dynamics enable bidirectional movement of APC-RNPs. **(a)** Kymographs showing full-length APC-647 (400 pM) tracking growing and shrinking microtubule plus ends decorated with EB1-mGFP (5000 pM) and unlabelled EB1 (5000 pM). Yellow box indicates zoomed part. Arrowheads highlight APC-647 on shrinking MT plus end. **(b)** Microtubule growth speed and extension periods in the presence and absence of APC-647 measured by tracking EB1-mGFP. **(c)** Kymographs showing RNP complexes of APC-647 (400 pM) and TMR-β2Btub_wt_ (12000 pM) tracking growing and shrinking microtubule plus ends decorated with EB1-mGFP (5000 pM) and unlabelled EB1 (5000 pM). Yellow box indicates zoomed part. Arrowheads highlight APC-647 and TMR-β2Btub_wt_ on shrinking MT plus end. **(d)** Microtubule growth speed and extension periods in the presence and absence of an APC-647 -TMR-β2Btub_wt_ complex measured by tracking EB1-mGFP. **(f)** 1-cumulative frequency plot showing dwell time of APC-647 (50 pM) on paclitaxel-stabilised microtubules dependent on EB1 concentration. Experimental data were fitted with a two-phase decay model. Inset: Slow and fast half-lives of dwell times were plotted against EB1 concentrations. **(g & h)** Kymographs showing processive RNP complexes of APC-647 (400 pM) and TMR-β2Btub_wt_ (12000 pM) in the presence of KIF3ABK (4000 pM), EB1-mGFP (5000 pM) and unlabelled EB1 (5000 pM). Three distinct stages are depicted: (i) processive transport of RNPs towards the growing MT plus end (arrows in (g & h)), (ii) growing plus-end-tracking of RNPs in the presence of EB1-mGFP (g & h), (iii) shrinking plus-end-tracking of RNPs in the absence of EB1-mGFP (arrowheads in (g)).

## Discussion

In this work, we show that APC-RNPs can be transported not only by the heterotrimeric kinesin-2 KIF3ABK, as previously shown (Baumann et al. 2020), but also by the homodimeric kinesin-1 KIF5AA. Compared to KIF3ABK, however, KIF5AA is less specific with weaker binding activities also to the β2Btub RNA. These weaker interactions of KIF5A with mRNP components are one possible cause of the often-observed remaining transport activity which persists after a key mRNA transport factor has been disabled (Dictenberg et al., 2008; Sharangdhar et al., 2017; Yoon et al., 2016).

We further show that APC is a versatile adaptor that specifically binds several G-rich motifs of localised neuronal mRNAs with high affinity and low off rates. Of note, FMRP binds the same motifs as APC, partially with higher affinities, which suggests a competition for the same mRNA targets, likely being controlled by relative subcellular protein abundance and PTMs in cells. For instance, phosphorylation of APC by GSK3β modulates the APC-MT interaction (Zumbrunn et al., 2001). Given that the MT- and RNA-binding regions overlap, a similar mechanism for APC-RNA binding might exist.

While KIF3ABK binds the ARM region of APC (Fig.S1a) via KAP3, KIF5A was reported to recognise a C-terminal APC fragment (Ruane et al., 2016 & Fig.S1a). This caused us to speculate that both kinesins can bind APC simultaneously, potentially increasing APC-RNP processivity. However, we could not detect any synergistic effect on APC transport parameters at limiting APC concentrations and in the presence of both kinesins. We rather find that all motility parameters except for the confinement ratio of APC in the presence of KIF3ABK and KIF5AA (scenario 2) are in between the parameters of the single motor conditions (scenarios 1 and 3). Specifically, the bimodal velocity distribution in the mixed motor condition argues for the presence of two distinct APC-kinesin populations and the mutually exclusive binding of KIF3ABK and KIF5AA to APC. Both motors might compete for the APC-ARM region, but at the chosen motor concentrations both motors transport APC almost equally.

We observed that the velocity of KIF5AA-based transport of APC fragments, β2Btub RNA and FMRP is faster than the transport of full-length APC. As this happens at experimental conditions under which mostly a dimeric APC binds a single KIF3ABK, and the velocity of KIF5AA-APC complexes does not correlate with motor intensity, we exclude the possibility that motor oligomers cause the velocity difference. We instead propose that full-length APC and APC-RNPs might use a special transport mode in which APC remains in contact with the MT while the motor drags it along the lattice. This idea is also supported by the extended dwell time of KIF5AA-APC complexes compared to cargo-free KIF5AA, which argues for additional MT attachment sites provided by APC. Such a transport mechanism could have advantages for long-range transport; if the components of the transport complex were not “lost” freely diffusing in the cytoplasmic space, the diffusion of one or both components along the MT could facilitate assembly of a new transport complex.

We finally show that EB1 is sufficient for recruiting APC and APC-RNPs to growing MT ends in vitro and that APC and APC-RNPs can track shrinking MT ends. Due to the high dwell times of full-length APC on the MT lattice, the observed shrinking-end-tracking is very likely caused by a continuous movement of APC-RNPs and not a rapid turnover as for EB proteins on the structural cap of MT plus ends (Bieling et al., 2007; Maurer et al., 2012). Since APC localises to MT plus ends within filopodia of axonal growth cones (Preitner et al., 2014), the combined action of active transport and growing- and shrinking-MT plus end tracking, could be an attractive mechanism to navigate bidirectionally within the growth cones and filopodia to scan for mRNP target sites.

As a microtubule plus-end-tracking protein, APC can further reach high local concentrations around growing MT ends, which could also contribute to a concentration-driven displacement of other proteins binding to similar mRNA motifs such as FMRP. As FMRP functions as a translational repressor (Darnell et al., 2011), such a plus-end-tracking-driven remodelling of mRNPs would be an interesting new mechanism to promote the translation of mRNAs from translationally inactive transport mRNPs upon their delivery to cellular domains with highly dynamic MTs, e.g. axonal growth cones. A plus-end-trailing of mRNPs has been observed in axonal growth cones (Leung et al., 2018) and it will now be of highest interest to understand which function interactions of mRNPs with growing MT ends could serve in the cell.

## Materials and Methods

### Materials

Chemicals were obtained from Sigma if not stated otherwise. Single-fluorophore, 5’-end-labelled RNA fragments from mouse β-actin mRNA (accession number: NM_007393.5) and mouse β2B-tubulin-mRNA (accession number: NM_023716.2) were purchased from IBA-Lifesciences (Germany). All 25 nt RNAs used for SPR experiments were purchased from IDT. The fragments for β-actin and β2Btub mRNAs were taken from the same transcripts listed above. The Map1b-fragment comes from accession number NM_008634.2, the Camk2α-mRNA fragment from NM_001286809.1. Total internal reflection fluorescence microscopy (TIRF-M) experiments were performed on an iMIC (TILL Photonics, Germany) TIRF microscope equipped with three Evolve 512 EMCCD cameras (Photometrics, UK), a 100x 1.49 NA objective lens (Olympus, Japan), a quadband filter (405/488/561/647, Semrock, USA) and four different laser lines (405 nm, 488 nm, 561 nm, 639 nm). An Olympus tube lens adds a post magnification of x1.33, which results in a final pixel size of 120.3 nm. SPR experiments were performed on a BIACORE T100 system (GE Healthcare) using the Twin-Strep-tag® Capture Kit (IBA-Lifesciences) and CM5-Chips (Cytiva).

## Methods

### Cloning and expression of recombinant proteins

#### APC-ARM, APC-basic & APC-C

##### Cloning

Cloning and expression of APC-ARM (amino acids 336-1016) was described previously (Baumann et al., 2020).

APC-basic (amino acids 2163-2670) and APC-C (amino acids 2163-2845) regions were PCR-amplified from the full-length APC (*Mus musculus* APC, accession no. AAB59632) coding sequence and inserted by Gibson assembly into a pCoofy17a (Scholz et al., 2013) backbone. Both genes were fused upstream with a Protein A and SUMO3 coding sequence and downstream with sequences for the HRV-3C protease cleavage site, a HL-linker (Baumann et al., 2020) and a Twin-Strep-tag®. Both constructs were validated by sequencing of the entire open-reading frame and named pCoofy17_ZZ-SUMO3-APCBasic-3C-HL-Strep and pCoofy17_ZZ-SUMO3-APCBasic/KIF5bs/EB1bs-3C-HL-Strep respectively.

##### Expression

Both constructs were chemically transformed into the Rosetta™(DE3) pLysS strain. The transformed cells were plated in LB agar containing 50 μg/μL kanamycin and 25 μg/μL chloramphenicol. After overnight incubation, a single colony of each clone was grown overnight in 10 mL of LB medium containing the mentioned antibiotics on a rotary shaker (200 rpm) at 37°C. On the next day, 200 mL of LB/antibiotics medium were inoculated with 10 mL of pre-culture medium and incubated under the same conditions. Once cells reached an OD600 of 3, 30 mL of the culture were used to inoculate 1 L of fresh LB/antibiotics medium. The cultures were incubated at 37°C while shaking until the cells reached the mid-exponential phase (OD600 ≃ 0.6), then induction of protein expression was performed by the addition of 1 mM IPTG and cells were incubated at 37°C while shaking for 3 h. Cells were then harvested by centrifugation at 7000 x g at 4°C, washed by resuspension with 1x PBS and harvested again by centrifugation at 7000 x g at 4°C. Cells were frozen in liquid N_2_ and stored at −80°C.

#### FMRP

##### Cloning

A vector containing the codon-optimised mouse FMR1 gene (accession no.: NM_008031.3) was linearised by PCR to remove the existing region containing the TEV protease cleavage site followed by a linker sequence (ILGAPSGGGATAGAGGAGGPA GLIN), a SNAP tag, the HRV-3C protease cleavage site and a Twin-Strep-tag®originally located at the C-terminal of the genes. The Twin-Strep-tag® was then added again at the C-terminus of each gene by Gibson assembly. The construct was validated by sequencing of the entire open-reading frame and named pLIB_ZZ-3C-FMRP-Strep.

##### Expression

600 mL Sf21 II cell suspension at 1.5×10e6 cells/mL were infected with 600 µL of a V1 Baculovirus and incubated in a 2 L flask in a shaker-incubator at 27ºC, 100 rpm for 3 days. Cells were harvested by centrifugation at 2000 x g for 15 min at 4ºC, resuspended in cold 1x PBS and centrifuged again at 2000 x g for 15 min at 4ºC. The supernatant was discarded and the pellet was frozen in liquid nitrogen and stored at −80ºC.

#### KIF5A

##### Cloning

KIF5A coding region was PCR-amplified from a codon-optimised version of mouse KIF5A (accession no.: NP_001034089.1) and inserted by Gibson assembly into a pLIB backbone vector for expression in insect cells. This resulted in a recombinant Kif5A gene flanked by a Protein A solubility tag followed by the HRV-3C protease cleavage site at the N-terminus and the TEV protease cleavage site, a linker sequence (ILGAPSGGGATAGAGGAGGPA GLIN), a SNAP tag, another HRV-3C protease cleavage site and a Twin-Strep-tag® at the C-terminus. The final construct was validated by sequencing of the entire open-reading frame and named pLIB-ZZ-3C-KIF5A-TEV-HL-SNAP-3C-Strep. *Expression:* 800 mL Sf21 II cell suspension at 1.5×10e6 cells/mL were infected with 800 µL of a V1 Baculovirus and incubated in a 2 L flask in a shaker-incubator at 27ºC, 100 rpm for 3 days. Cells were harvested by centrifugation at 2000 x g for 15 min at 4ºC, resuspended in cold 1x PBS and centrifuged again at 2000 x g for 15 min at 4ºC. The supernatant was discarded and the pellet was frozen in liquid nitrogen and stored at −80ºC.

#### KIF5B

##### Cloning

A plasmid containing the KIF5B coding sequence (accession no.: NM_008448.3) was obtained through MTA from Addgene (plasmid 127617). The KIF5B region was PCR-amplified from this plasmid and inserted by Gibson assembly into a backbone vector already existing in the lab stock harbouring a Protein A solubility tag followed by the HRV-3C protease cleavage site at the N-terminus and the TEV protease cleavage site, a linker sequence (HL-linker), a SNAP tag, another HRV-3C protease cleavage site and a Twin-Strep® purification tag at the C-terminus. The final construct was validated by sequencing of the entire open-reading frame and named pLIB-ZZ-3C-KIF5B-TEV-HL-SNAP-3C-Strep.

##### Expression

700 mL Sf21 III cell suspension at 1.5×10e6 cells/mL were infected with 2000 µL of a V1 Baculovirus and incubated in a 2 L flask in a shaker-incubator at 27ºC, 100rpm for 3 days. Cells were harvested by centrifugation at 2000 x g for 15 min at 4ºC, resuspended in cold 1x PBS and centrifuged again at 2000 x g for 15 min at 4ºC. The supernatant was discarded and the pellet was frozen in liquid nitrogen and stored at −80ºC.

#### MAP7

##### Cloning

The human MAP7 coding region (accession no.: NM_018067.5) and optionally a GSA-linker-SNAP-tag fragment were PCR-amplified and inserted by Gibson assembly into the backbone vector pLIB from the BiGBac System for insect cell-based protein expression. This resulted in two new vectors, in the first one the MAP7 gene has a Protein A solubility tag followed by the HRV-3C protease cleavage site at the N-terminal and another HRV-3C protease cleavage site and a Twin-Strep-tag® at the C-terminus. In the second vector, the MAP7 gene also has a SNAP tag at its C-terminus before the HRC-3C protease cleavage site. The final constructs were validated by sequencing of the entire open-reading frame and named pLIB_ZZ-3C-MAP7-3C-Strep and pLIB_ZZ-3C**-**MAP7-GSAlink-SNAP-3C-Strep.

##### Expression

600 mL Sf21 9 cell suspension at 2×10e6 cells/mL were infected with 600 µL of a V1 Baculovirus and incubated in a 2 L flask in a shaker-incubator at 27ºC, 100 rpm for 3 days. Cells were harvested by centrifugation at 2000 x g for 15 min at 4ºC, resuspended in cold 1x PBS and centrifuged again at 2000 x g for 15 min at 4ºC. The supernatant was discarded and the pellet was frozen in liquid nitrogen and stored at −80ºC EB1 and EB1-mGFP were expressed from previously described constructs and purified according to published protocols (Maurer et al., 2014).

#### Protein biochemistry

##### APC-ARM

APC-ARM was purified as described previously (Baumann et al., 2020).

##### APC-basic & APC-C purification

50 mL of cold purification buffer (80 mM NaPi pH 7.2, 800 mM KCl, 5 mM MgCl_2_, 5 mM DTT, 2.5 mM EDTA, 0.1% Tween-20) supplemented with 1 cOmplete™ ULTRA Tablet (Sigma) and DNase I (Roche) were added to 3 g of frozen cells obtained from 1 L of bacterial culture expressing the respective APC-basic construct. In the case of APC-C, 3 pellets of 3 g each were resuspended with 150 mL of the same purification buffer. The pellets were thawed at 4°C and resuspended by stirring. The bacteria were disrupted at a pressure of 5,500 kPa with a cell disruptor (Emulsiflex, Avestin), the lysates were clarified by centrifugation (184,000 x g, 45 min, 4°C) and the supernatants were applied to distinct 5 mL StrepTrap HP columns (Cytiva). The columns were first washed with 40 column volumes (CV) of purification buffer, then with 20 CV of wash buffer A (80 mM NaPi pH 7.2, 500 mM KCl, 5 mM MgCl_2_, 5 mM DTT, 2.5 mM EDTA, 0.1% Tween-20) and finally with 20 CVs of wash buffer B (80 mM NaPi pH 7.2, 300 mM KCl, 5 mM MgCl_2_, 5 mM DTT, 2.5 mM EDTA, 0.1% Tween-20). The salt concentration in the wash buffers was slowly reduced to avoid protein precipitation. APC-basic and APC-C were eluted from the columns by flowing 5 CV of elution buffer containing 80 mM NaPi, 300 mM KCl, 5 mM MgCl_2_, 2.5 mM DTT, 2.5 mM EDTA and 10 mM desthiobiotin through them. SUMO3 tags were removed by incubating the eluted proteins overnight at 4°C with SENP2 protease. On the next day, the proteins were concentrated using Amicon® Ultra 15 mL Centrifugal Filters (Merck), ultra-centrifuged (280,000 x g, 10 min, 4°C) and gel-filtered using a Superdex 200 10/300 GL column (Cytiva). Peak fractions were pooled according to the level of purity shown by SDS-PAGE analysis and the concentration of the pure proteins was measured by Bradford assay. APC-C and APC-basic were aliquoted on ice and snap frozen in liquid N_2_.

##### FMRP purification

50 mL of RT purification buffer (50 mM HEPES-NaOH pH 8, 50 mM glutamate, 50 mM arginine, 250 mM glucose, 500 mM KCl, 5 mM MgCl_2_, 2.5 mM DTT, 1 mM EDTA, 0.001% Brij-35 and 10 mM ATP) supplemented with 1 cOmplete™ ULTRA Tablet (Sigma) and DNase I (Roche) were added to 3 g of frozen cells obtained from 600 mL of Sf21 insect cell culture expressing the respective FMRP construct. The pellets were thawed at RT and resuspended by stirring. The cells were disrupted with a Dounce homogeniser, the lysate was clarified by centrifugation (184,000 x g, 45 min, 20°C) and the supernatant was applied to a 5 mL StrepTrap HP column (Cytiva) at RT. The column was washed with 40 CV of purification buffer and FMRP was eluted from the column by flowing 3 CV of elution buffer containing 50 mM HEPES-NaOH pH 8, 50 mM glutamine, 50 mM arginine, 250 mM glucose, 500 mM KCl, 5 mM MgCl_2_, 2.5 mM DTT, 1 mM EDTA, 0.001% Brij-35, 10 mM ATP and 30 mM desthiobiotin through the column. FMRP was concentrated using Amicon® Ultra 15 mL Centrifugal Filters (Merck) and desthiobiotin was removed with a PD-10 Desalting Column (Cytiva). The level of purity was assessed by SDS-PAGE analysis and the concentration of the pure protein was measured by Bradford assay. FMRP was aliquoted on ice and snap frozen in liquid N_2_.

##### KIF5A purification

50 mL of cold purification buffer (80 mM NaPi pH 7.2, 400 mM KCl, 5 mM MgCl_2_, 2.5 mM DTT, 2.5 mM EDTA, 0.001% Brij-35 and 0.1 mM ATP) supplemented with 1 cOmplete™ ULTRA Tablet (Sigma) and DNase I (Roche) were added to 3 g of frozen cells obtained from 800 mL of Sf21 insect cell culture expressing the respective KIF5A construct. The pellet was thawed at RT and resuspended by stirring. The cells were disrupted with a Dounce homogeniser, the lysate was clarified by centrifugation (184,000 x g, 45 min, 4°C) and the supernatant was applied to a 5 mL StrepTrap HP column (Cytiva). The column was washed with 40 CV of purification buffer and KIF5A was eluted from the column by flowing 3 CV of elution buffer containing 50 mM HEPES-NaOH pH 8, 50 mM glutamine, 50 mM arginine, 250 mM glucose, 500 mM KCl, 5 mM MgCl_2_, 2.5 mM DTT, 1 mM EDTA, 0.001% Brij-35, 10 mM ATP and 15 mM desthiobiotin through the column. KIF5A was concentrated using Amicon® Ultra 15 mL Centrifugal Filters (Merck) and desthiobiotin was removed with a PD-10 Desalting Column (Cytiva). The eluted protein was split in two aliquots, one was incubated with His-tagged Prescission protease and His-tagged TEV protease at 4°C overnight to cleave off solubility and affinity tags respectively, and the other aliquot was incubated with His-tagged Prescission protease only to remove just the solubility tags and maintain the SNAP tag for subsequent SNAP tag labelling SNAP-reactive dyes. On the next day, both fully cleaved KIF5A and KIF5A-SNAP were passed through a 1 mL StrepTrap HP column (Cytiva) to remove potentially uncleaved protein. KIF5A-SNAP was incubated with Alexa Fluor 647 dye (New England Bioscience) at a ratio of 1:2 protein to dye for 2 hours at 15°C, then applied onto a Zeba™ Spin desalting columns (Thermo Fisher Scientific) to remove the excess of dye. Both proteins, fully cleaved KIF5A and Alexa647-labelled KIF5A were then concentrated using Amicon® Ultra 15 mL Centrifugal Filters (Merck), ultra-centrifuged (280,000 x g, 10 min, 4°C) and gel-filtered using a Superose6 10/300 GL column (GE Healthcare). Peak fractions were pooled according to the level of purity shown by SDS-PAGE analysis and the concentration of the pure proteins was measured by Bradford assay. KIF5A and 647-KIF5A were aliquoted on ice and snap frozen in liquid N_2_.

##### KIF5B purification

200 mL of cold lysis buffer (80 mM HEPES-NaOH pH 7.4, 50 mM glutamate, 50 mM arginine, 500 mM NaCl, 1 mM MgCl_2_, 0.5% Triton-X100, 2 mM DTT, 1 mM EGTA, 0.1 mM ATP and 10% glycerol) supplemented with 4 cOmplete™ ULTRA Tablet (Sigma) and DNase I (Roche) were added to 20 g of frozen cells obtained from 700 mL of an Sf21 insect cell culture expressing the respective KIF5B construct. The pellets were thawed on ice and resuspended by stirring. The cells were lysed using a French press at a pressure of 2,500 psi, the lysate was clarified by centrifugation (184,000 x g, 45 min, 4°C) and the supernatant was applied to 2x 5 mL StrepTrap HP columns (Cytiva) at 4°C. The columns were first washed with 40 CV of lysis buffer and subsequently with 40 CV of wash buffer (80 mM HEPES-NaOH pH 7.4, 50 mM glutamate, 50 mM arginine, 300 mM NaCl, 1 mM MgCl_2_, 0.05% Triton-X100, 1 mM DTT, 1 mM EGTA, 0.1 mM ATP and 10% glycerol). The Twin-Strep-tag® was removed by on-column cleavage with His-tagged TEV protease overnight at 4°C. KIF5B was eluted with 20 CV of wash buffer, concentrated using Amicon® Ultra 15 mL Centrifugal Filters (Merck), ultra-centrifuged (280,000 x g, 10 min, 4°C) and gel-filtered using a Superose 6 10/300 GL column (Cytiva) equilibrated with GF buffer (80 mM HEPES-NaOH pH 7.4, 50 mM glutamate, 50 mM arginine, 150 mM NaCl, 1 mM MgCl_2_, 1 mM DTT and 0.1 mM ATP). Peak fractions were pooled according to the level of purity shown by SDS-PAGE analysis and the concentration of the pure protein was measured by Bradford assay. KIF5B was aliquoted on ice and snap frozen in liquid N_2_.

##### MAP7 purification

50 mL of cold lysis buffer (80 mM sodium phosphate pH 7.2, 50 mM glutamate, 50 mM arginine, 500 mM KCl, 1 mM MgCl_2_, 0.001% Brij-35, 1 mM DTT, 1 mM EDTA and 0.1 mM ATP) supplemented with 1 cOmplete™ ULTRA Tablet (Sigma) and DNase I (Roche) were added to 5 g of frozen cells obtained from 600 mL of Sf21 insect cell culture expressing the MAP7 construct. The pellets were thawed at 4°C and resuspended by stirring. The cells were lysed using a French press at a pressure of 2,500 psi, the lysate was clarified by centrifugation (184,000 x g, 45 min, 4°C) and the supernatant was applied to a 5 mL StrepTrap HP column (Cytiva) at 4°C. The column was washed with 30 CV of lysis buffer, the protein was eluted with 3 CV of elution buffer (80 mM sodium phosphate, 50 mM glutamate, 50 mM arginine, 500 mM KCl, 1 mM MgCl_2_, 0.001% Brij-35, 1 mM DTT, 1 mM EDTA, 0.1 mM ATP and 5 mM desthiobiotin) and concentrated using Amicon® Ultra 15 mL Centrifugal Filters (Merck). Desthiobiotin was removed using a PD-10 Desalting Column (Cytiva) and the protein was gel-filtered using a Superdex200 Increase 10/300 GL column (Cytiva) equilibrated with lysis buffer. After solubility and affinity tags were removed using His-tagged 3C PreScission protease, MAP7 was ultra-centrifuged (280,000 x g, 10 min, 4°C) and then again gel-filtered using a Superdex200 Increase 10/300 GL column (Cytiva) equilibrated with lysis buffer. Peak fractions were concentrated using Amicon® Ultra Centrifugal Filters (Merck) and the concentration of the pure protein was measured by Bradford assay. MAP7 was aliquoted on ice and snap frozen in liquid N_2_.

##### Purification of EB1 & EB1-mGFP

50 mL of cold lysis buffer (50 mM sodium phosphate pH 7.2, 400 mM KCl, 2 mM MgCl_2_ x 6 H_2_O, 0.002% Brij35, 1 mM β-mercaptoethanol, 1 mM imidazole) supplemented with### cOmplete™ ULTRA Tablet (Sigma) and DNase I (Roche) were added to frozen cell pellets expressing the respective EB1 and EB1-mGFP constructs, respectively. Pellets were thawed and sonicated. The lysates were clarified by centrifugation (184,000 x g, 45 min, 4°C) and the supernatants were applied to 5 mL HiTrap columns (GE Healthcare) loaded with cobalt(II)-chloride hexahydrate (Sigma). The columns were washed with EB1 wash buffer A (50 mM sodium phosphate pH 7.2, 400 mM KCl, 2 mM MgCl_2_ x 6 H_2_O, 0.002% Brij35, 1 mM β-mercaptoethanol, 10 mM imidazole) and the proteins were eluted with EB1 elution buffer B (50 mM sodium phosphate pH 7.2, 400 mM KCl, 2 mM MgCl_2_ x 6 H_2_O, 0.002% Brij35, 1 mM β-mercaptoethanol, 500 mM imidazole). The eluate was concentrated with a Vivaspin concentrator (Sartorius) and supplemented with 8 mM β-mercaptoethanol. Affinity tags were cleaved off overnight at 4°C using TEV protease. Gel filtration was performed using a Superdex200 10/300 GL column (Cytiva). Peak fractions were pooled according to the level of purity shown by SDS-PAGE analysis and the concentrations of the pure proteins were measured by Bradford assay. EB1 and EB1-mGFP were aliquoted on ice and snap frozen in liquid N_2_.

#### SNAP labelling of proteins

SNAP labelling was carried out at 4°C for 16 hours using 2-fold access of dye over protein. Fresh DTT was added to a concentration of 2 mM directly before labelling. Unbound dye was immediately removed using Zeba™ Spin Desalting Columns (Thermo Fisher Scientific), and subsequently the labelled protein was processed by size exclusion chromatography to remove potential aggregates.

#### SDS PAGE and Western blotting

SDS-PAGE was performed using the Mini-Protean system and TGX precast gels (Biorad). Proteins were stained with BlueSafe solution (NZYtech). Western blotting on SDS gels was performed with an Invitrogen iBlot2 device and fitting iBlot2 NC stacks (Invitrogen). To detect protein expression, antibodies against the Twin-Strep-tag® (Strep-HRP tag II Novagen 71591-3 dilution 1/4000) and the Protein A sequence (ProtA-HRP Abcam ab18596 dilution 1/10000) were used.

#### Reproducibility statistical analysis

At least three independent experiments were performed for all tested conditions with the exception of experiments analysed in: Fig.4f – one experiment for the condition with diffusive APC-647 (brown curve), Fig.4h – one experiment for the condition KIF5AA + full-length APC-647, Fig.6f – one experiment for each EB1 concentration.

#### Statistical analysis

Statistical analysis was performed with Graphpad Prism and Origin. The types of tatistical tests performed are indicated in figure legends. Statistical significance in figures is indicated by * p < 0.05, ** p < 0.01, *** p < 0.001 and **** p < 0.0001. In case no significant difference between median or mean of samples was detected, the p value is indicated.

#### Polymerisation and stabilisation of MTs

Microtubules were polymerised from bovine tubulin with stochastic incorporation of ATTO390 (Attotech, Germany)-and biotin (ThermoFisher, #20217)-labelled tubulin. Mixes of 36 μM unlabelled tubulin, 14 μM ATTO390-tubulin, and 10 μM biotin-tubulin were incubated in BRB80 (80 mM K-PIPES pH 6.85, 2 mM MgCl_2_, 1 mM EGTA) with 4 mM GTP (Sigma) for 25 min at 37°C in a total volume of 50 uL. Next, pre-warmed BRB80 supplemented with 20 µM paclitaxel (Sigma) was added and incubation continued for 60 min at 37°C. Polymerised microtubules were pelleted at 20,000 x g for 5 min and resuspended in 60 µL BRB80 supplemented with 20 µM paclitaxel. For the final assay, MTs are diluted 1:8 in RT assay buffer (60 µL volume) before flowing them into the assay chamber.

#### Polymerisation of MT seeds

MT seeds were polymerised from bovine tubulin by incubating 20 μM unlabelled tubulin, 10 μM ATTO390-tubulin and 9 μM biotin-tubulin in BRB80 (80 mM K-PIPES pH 6.85, 2 mM MgCl_2_, 1 mM EGTA) with 0.5 mM GMPCPP (Jena Bioscience) for 17 min at 37°C in a total volume of 40 uL. Next, 400 µL pre warmed BRB80 were added and polymerised seeds were pelleted at 20,000 x g for 7 min and resuspended in 50 µL BRB80. For the final assay, seeds are diluted 1:5 in RT BRB80 and then 1:12 in RT assay buffer (60 µL volume) before flowing into the assay chamber.

#### Motility chamber preparation

Glass surfaces were prepared as described previously. Motility chambers with a volume of ∼10 μL were assembled by adhering glass cover slips functionalised with PEG and biotin-PEG (Microsurfaces Inc, USA, #Bio_01(2007134-01)) to glass slides passivated with PLL(20)-g[3.5]-PEG(2) (SuSos AG, Duebendorf, Switzerland) using 2 parallel segments of double-sided adhesive tape. Chamber surfaces were passivated for 5 min on ice with 5% (w/v) Pluronic F-127 #P2443) in TIRF-M assay buffer (for paclitaxel-stabilised MTs: 90 mM HEPES, 10 mM PIPES #P6757, 2.5 mM MgCl_2_, 1.5 mM EGTA #03777, 15 mM β-mercaptoethanol, pH 6.95. For dynamic MTs: 45 mM HEPES, 5 mM PIPES #P6757, 5 mM MgAc, 1.5 mM EGTA #03777, 15 mM β-mercaptoethanol, pH 6.9; note, that the buffer was adjusted to compensate for 11.8 mM BRB80 that is introduced into the final assay as storage buffer of free tubulin) supplemented with 50 ug/mL kappa-casein #C0406. Chambers were then flushed with 25 ug/mL NeutrAvidin (Invitrogen, #A-2666) in assay buffer containing 200 ug/mL kappa-casein and instantly rinsed with assay buffer followed by warm up to RT. After 3 min stabilised MTs or seeds were diluted in RT assay buffer and added to the chamber followed by 3 min incubation.

#### In vitro motility assay

Constituents of motility assays were incubated on ice for 15 ± 1 min in TIRF-M assay buffer supplemented with 4% PEG-3350 #1546547, 2.5 mM ATP and 0.5 U/µL SuperaseIn (Invitrogen, AM2694). Pre incubation mixes typically contained (a) MT motor proteins: 35 nM KIF3ABK, 10.5 nM KIF5AA, 10.5 nM KIF5BB; (b) RBPs: 7 nM APC, 56 nM FMRP, 56 nM PURα; (c) RNAs: 140 nM RNA. The pre incubation mix was diluted 69.89-fold into final motility buffer lacking RNase inhibitor and containing additionally 0.32 mg/mL glucose oxidase (AMSBIO, Germany, #22778.02), 0.275 mg/mL catalase #C40, 50 mM glucose #G7021, 50 µg/mL beta-casein #C6905 and 0.12% methylcellulose #M0512. Exact final protein and RNA concentrations are given in figure legends. Diluted mixes were added to pre immobilised microtubules in the motility chamber for imaging at 30 ± 1°C.

#### In vitro dynamic MT assay

Constituents of dynamic assays were incubated on ice for 18 ± 1 min in TIRF-M assay buffer supplemented with 2% PEG-3350 #1546547, 2 mM ATP and 0.5 U/µL SuperaseIn (Invitrogen, AM2694). Typical protein concentrations were: 28 nM APC and 840 nM RNA. The preincubation mix was diluted 69.89-fold into final motility buffer lacking RNase inhibitor and containing additionally 5 µM EB1, 5 µM EB1-mGFP, 25 µM tubulin, 1 mM GTP, 0.32 mg/mL glucose oxidase (AMSBIO, Germany, #22778.02), 0.275 mg/mL catalase #C40, 50 mM glucose #G7021, 50 ug/mL beta-casein #C6905 and 0.12% methylcellulose #M0512. Exact final protein and RNA concentrations are given in figure legends. Diluted mixes were added to pre immobilised microtubule seeds in the motility chamber for imaging at 30 ± 1°C.

#### TIRF microscopy

For triple colour imaging on stabilised MTs two channels were recorded alternating with 4 frames per second (fps, 125 ms exposure per channel) by switching laser lines 561 nm and 639 nm. ATTO390 microtubules were recorded as a single image after time lapse movies had been finalised. This protocol was chosen since ATTO390 bleaches comparably fast and total illumination time per cycle is saved to increase the frame rate in the other channels. For triple colour imaging with MTs three channels were recorded alternating, with 1.33 or 2.22 fps (250 or 150 ms exposure per channel) by switching laser lines 488 nm, 561 nm and 639 nm. Laser powers as well as exposure times and acquisition frame rates were kept constant within a set of experiments. For channel alignments, images with 100 µm TetraSpeck fluorescent beads (Invitrogen, UK) were recorded in all channels before experiments were started.

#### Analysis of TIRF-M data

For analysis of dynamic properties and fluorescence intensities of APC-RNA complexes (APC-RNPs), TIRF-M movies were loaded into the FIJI software and analysed using TrackMate (Baumann et al., 2020; Grawenhoff et al., 2022; Schindelin et al., 2012; Tinevez et al., 2017). In all cases, entire TIRF-M movies including the entire field of view were analysed to guarantee an unbiased analysis. Single particles were detected using the DoG (Difference of Gaussian) detector with an estimated spot diameter of 800 nm. Intensity thresholds for spot detection depended on the amount of fluorescent species used in the respective experiments and was kept constant for all comparisons of experiments using the same concentrations of fluorescent proteins or RNAs and the same imaging settings. To compute trajectories of diffusive particles, we used the “Simple LAP tracker” with the following settings: maximal linking distance = 600 nm, maximal gap-closing distance = 600 nm, maximal frame gap = 2. To exclude stationary and long-range particles as well as background signals, only trajectories with track displacement > 1000 nm and maximum distance < 7000 nm were included. For the analysis of processive RNPs we used the “Linear motion LAP tracker” with the following settings: initial search radius = 600 nm, search radius = 600 nm, maximal frame gap = 2. To exclude stationary and diffusive particles as well as background signals, only trajectories with maximum distance < 6000 nm or < 7000 nm (Fig.4) were included. For tracking of EB1-GFP at dynamic MT plus ends, we used the “Simple LAP tracker” with the following settings: maximal linking distance = 1000 nm, maximal gap-closing distance = 1000 nm, maximal frame gap = 2. Filters for spots in track > 100 and track displacement > 1000 nm were applied. Setting the threshold in TrackMate depends on wavelength, laser power and fluorophore concentration. Typical values for mGFP, TMR and Alexa647 were 35, 25 and 10, respectively. The background fluorescence of TIRF-M data was measured by calculating the mean intensity in a 28 × 18-pixel rectangular box at five different positions (upper left & right, center, lower left & right) of the field of view at four different time points of the movie. The obtained mean background fluorescence was subtracted from total spot intensities obtained by Trackmate single-particle tracking.

#### Analysis of RNP dynamics

To analyse dwell times of transport RNP components, 1-cumulative frequency distributions of the respective dwell times (=track duration) were fitted to the monoexponential or bioexponential decay functions *y* = *y*_0_ + *Ae*^−*x/t*^ and *y* = *y*_0_ + *A*_1_*e*^−*x/t1*^ + *A*_2_*e*^−*x/t2*^. The decay constant τ was derived by τ = *t* * ln (2). The mean velocity of RNA transport complexes was determined by computing the average of individual mean track velocities. The mean track velocity is the mean of the instantaneous velocities of a track. To determine the confinement ratio of RNP components the net displacement of a particle is divided by its total distance travelled.

#### Surface Plasmon Resonance

Surface Plasmon Resonance (SPR) experiments were performed with a BIACORE T100 system (GE Healthcare). Strep-Tactin XT® from Twin-Strep tag® Capture Kit (IBA-Lifesciences) was covalently amine-coupled to a CM5-Chip (Cytiva) in both working and reference channel according to the manufacturer’s instructions. The proteins APC-basic and APC-C were diluted to a working concentration of 1 ug/mL in HBS-P buffer (10 mM HEPES pH 7.4, 150 mM NaCl, 0.05% surfactant P20) and captured on the working channel only through the interaction between their Twin-Strep-tag® and the Strep-Tactin XT® previously immobilised to the chip. Five different RNAs were diluted in the same HBS-P buffer by serial dilution of 1:2 from 500 nM to 0.49 nM for a total of 11 different concentrations. Kinetic analysis of protein–RNA interaction was performed at a flow rate of 30 μL per min in HBS-P buffer. To assess protein-RNA interaction each concentration of RNA was injected in both reference and working channel for 5 minutes, subsequent dissociation was allowed for 5 min in HBS-P buffer. To remove any residual bound protein and RNA, 1 min regeneration injection with 3 M GuHCl was performed in both working and reference channels. To calculate the average values for the analyte response at equilibrium the binding curves were referenced against the signal from the ligand-free reference channel and 3 buffer runs. Kinetic fits and KD calculations were performed with the BIAevaluation software (GE Healthcare) using the two-state binding model available in the software. Steady-state binding curves were obtained by plotting the response at equilibrium against analyte concentration and curve fitting with the OriginPro2015 software using OneSiteBind 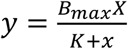 and TwoSiteBind 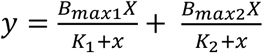 models. Every RNA concentration was tested in duplicates within each experiment and all experiments were performed in duplicate on different days, the results of the 2 experiments were averaged.

## Acknowledgments

We thank the CRG BMS-PT core facility for support by providing reagents and training on instrumentation.

## Author contributions

SJB and SPM conceived the project. SJB, JG, ECR, SS, MG and AK performed experiments. SJB, JG, SPM and SS analysed the data. SPM, SJB and JG wrote the article. SPM supervised the project.

## Data and materials availability

All data needed to evaluate the conclusions in the paper are present in the paper and/or the Supplementary Materials. Additional data related to this paper may be requested from the authors.

## Supplementary Materials

Figure S1 relates to Fig.1

Figure S2 related to Fig.2

Figure S3 relates to Fig.3

Figure S4 relates to Fig.6

**Movie S1**. KIF3ABK transports APC–β2B-tubulin RNA complexes (top) and KIF5AA. transports APC–β2B-tubulin RNA complexes (middle) or β2B-tubulin RNA (bottom).

**Movie S2**. KIF5AA transports APC-mGFP.

**Movie S3**. KIF5AA transports mGFP-FMRP.

**Movie S4**. KIF5AA-647 (magenta) transports APC-TMR (cyan).

**Movie S5**. EB1 (cyan)-dependent plus-end-tracking of APC (red). Channels are shifted slightly to allow better assessment of the individual behaviour of both proteins.

**Movie S6**. Tracking of EB1-mGFP using the Trackmate plugin for Fiji. Raw movie (top), particle detection (middle), track overlay (bottom).

**Movie S7**. Plus-end-tracking of APC (red) – β2B-tubulin RNA (yellow) complexes. Channels are shifted slightly to allow better assessment of the individual behaviour of both proteins.

**Movie S8**. Plus-end-tracking of EB1 (left), APC (middle) and β2B-tubulin RNA (right).

**Movie S9**. APC (middle) and β2B-tubulin RNA (right) on dynamic MTs in the absence of EB1.

**Movie S10**. EB1 (left), β2B-tubulin RNA (right) on dynamic MTs in the absence of APC.

**Movie S11**. APC–β2B-tubulin RNA complexes (yellow) are transported by KIF3ABK and plus-end-track via EB1 (cyan) on dynamic MTs. The white lines serve as guides to follow one RNA throughout a complex trajectory of growing & shrinking end tracking and active KIF3ABK-based transport.

## References

Akhmanova, A., and Steinmetz, M.O. (2008). Tracking the ends: a dynamic protein network controls the fate of microtubule tips. Nat. Rev. Mol. Cell Biol. 9, 309–322.

Batish, M., van den Bogaard, P., Kramer, F.R., and Tyagi, S. (2012). Neuronal mRNAs travel singly into dendrites. Proc. Natl. Acad. Sci. 109, 4645–4650.

Baumann, S., Komissarov, A., Gili, M., Ruprecht, V., Wieser, S., and Maurer, S.P. (2020). A reconstituted mammalian APC-kinesin complex selectively transports defined packages of axonal mRNAs. Sci. Adv. 6, eaaz1588.

Bieling, P., Laan, L., Schek, H., Munteanu, E.L., Sandblad, L., Dogterom, M., Brunner, D., and Surrey, T. (2007). Reconstitution of a microtubule plus-end tracking system in vitro. Nature 450, 1100–1105.

Bieling, P., Telley, I.A., Hentrich, C., Piehler, J., and Surrey, T. (2010). Fluorescence Microscopy Assays on Chemically Functionalized Surfaces for Quantitative Imaging of Microtubule, Motor, and +TIP Dynamics. In Methods in Cell Biology, (Elsevier), pp. 555– 580.

Buxbaum, A.R., Haimovich, G., and Singer, R.H. (2015). In the right place at the right time: visualizing and understanding mRNA localization. Nat. Rev. Mol. Cell Biol. 16, 95–109.

Coles, C.H., and Bradke, F. (2014). Microtubule self-organization via protein-RNA network crosstalk. Cell 158, 245–247.

Darnell, J.C., Jensen, K.B., Jin, P., Brown, V., Warren, S.T., and Darnell, R.B. (2001). Fragile X mental retardation protein targets G quartet mRNAs important for neuronal function. Cell 107, 489–499.

Darnell, J.C., Van Driesche, S.J., Zhang, C., Hung, K.Y.S., Mele, A., Fraser, C.E., Stone, E.F., Chen, C., Fak, J.J., Chi, S.W., et al. (2011). FMRP Stalls Ribosomal Translocation on mRNAs Linked to Synaptic Function and Autism. Cell 146, 247–261.

Dictenberg, J.B., Swanger, S.A., Antar, L.N., Singer, R.H., and Bassell, G.J. (2008). A Direct Role for FMRP in Activity-Dependent Dendritic mRNA Transport Links Filopodial-Spine Morphogenesis to Fragile X Syndrome. Dev. Cell 14, 926–939.

Dimitrova-Paternoga, L., Jagtap, P.K.A., Cyrklaff, A., Vaishali Lapouge, K., Sehr, P., Perez, K., Heber, S., Löw, C., Hennig, J., et al. (2021). Molecular basis of mRNA transport by a kinesin-1–atypical tropomyosin complex. Genes Dev. 35, 976–991.

Feierbach, B., Verde, F., and Chang, F. (2004). Regulation of a formin complex by the microtubule plus end protein tea1p. J. Cell Biol. 165, 697–707.

Ferro, L.S., Fang, Q., Eshun-Wilson, L., Fernandes, J., Jack, A., Farrell, D.P., Golcuk, M., Huijben, T., Costa, K., Gur, M., et al. (2022). Structural and functional insight into regulation of kinesin-1 by microtubule-associated protein MAP7. Science (80-.). 375, 326–331.

Goering, R., Hudish, L.I., Guzman, B.B., Raj, N., Bassell, G.J., Russ, H.A., Dominguez, D., and Taliaferro, J.M. (2020). FMRP promotes RNA localization to neuronal projections through interactions between its RGG domain and G-quadruplex RNA sequences. Elife 9, 784728.

Grawenhoff, J., Baumann, S., and Maurer, S.P. (2022). In Vitro Reconstitution of Kinesin-Based, Axonal mRNA Transport. Methods Mol. Biol. 2431, 547–568.

Hooikaas, P.J., Martin, M., Mühlethaler, T., Kuijntjes, G.-J., Peeters, C.A.E., Katrukha, E.A., Ferrari, L., Stucchi, R., Verhagen, D.G.F., van Riel, W.E., et al. (2019). MAP7 family proteins regulate kinesin-1 recruitment and activation. J. Cell Biol. 218, 1298–1318.

Janke, C., and Magiera, M.M. (2020). The tubulin code and its role in controlling microtubule properties and functions. Nat. Rev. Mol. Cell Biol. 21, 307–326.

Jiang, K., Toedt, G., Montenegro Gouveia, S., Davey, N.E., Hua, S., Van Der Vaart, B., Grigoriev, I., Larsen, J., Pedersen, L.B., Bezstarosti, K., et al. (2012). A proteome-wide screen for mammalian SxIP motif-containing microtubule plus-end tracking proteins. Curr. Biol. 22, 1800–1807.

Jimbo, T., Kawasaki, Y., Koyama, R., Sato, R., Takada, S., Haraguchi, K., and Akiyama, T. (2002). Identification of a link between the tumour suppressor APC and the kinesin superfamily. Nat. Cell Biol. 4, 323–327.

Kanai, Y., Dohmae, N., and Hirokawa, N. (2004). Kinesin Transports RNA. Neuron 43, 513– 525.

Kapitein, L.C., and Hoogenraad, C.C. (2015). Building the Neuronal Microtubule Cytoskeleton. Neuron 87, 492–506.

Kislauskis, E.H., Zhu, X., and Singer, R.H. (1997). beta-Actin messenger RNA localization and protein synthesis augment cell motility. J. Cell Biol. 136, 1263–1270.

Leung, K.-M., Lu, B., Wong, H.H.-W., Lin, J.Q., Turner-Bridger, B., and Holt, C.E. (2018). Cue-Polarized Transport of β-actin mRNA Depends on 3′UTR and Microtubules in Live Growth Cones. Front. Cell. Neurosci. 12.

Litman, P., Barg, J., Rindzoonski, L., and Ginzburg, I. (1993). Subcellular localization of tau mRNA in differentiating neuronal cell culture: implications for neuronal polarity. Neuron 10, 627–638.

Martin, S.G., and Chang, F. (2003). Cell Polarity : A New Mod (e) of Anchoring Microtubules play a central role in the establishment. Curr. Biol. 13, 711–713.

Maurer, S.P., Bieling, P., Cope, J., Hoenger, A., and Surrey, T. (2011). GTPgammaS microtubules mimic the growing microtubule end structure recognized by end-binding proteins (EBs). Proc. Natl. Acad. Sci. U. S. A. 108, 3988–3993.

Maurer, S.P., Fourniol, F.J., Bohner, G., Moores, C.A., and Surrey, T. (2012). EBs Recognize a Nucleotide-Dependent Structural Cap at Growing Microtubule Ends. Cell 149, 371–382.

Maurer, S.P., Cade, N.I., Bohner, G., Gustafsson, N., Boutant, E., and Surrey, T. (2014). EB1 accelerates two conformational transitions important for microtubule maturation and dynamics. Curr. Biol. 24, 372–384.

Meijering, E., Dzyubachyk, O., and Smal, I. (2012). Methods for cell and particle tracking. Methods Enzymol. 504, 183–200.

Messitt, T.J., Gagnon, J.A., Kreiling, J.A., Pratt, C.A., Yoon, Y.J., and Mowry, K.L. (2008). Multiple Kinesin Motors Coordinate Cytoplasmic RNA Transport on a Subpopulation of Microtubules in Xenopus Oocytes. Dev. Cell 15, 426–436.

Mili, S., Moissoglu, K., and Macara, I.G. (2008). Genome-wide screen reveals APC-associated RNAs enriched in cell protrusions. Nature 453, 115–119.

Mimori-Kiyosue, Y., Shiina, N., and Tsukita, S. (2000). The dynamic behavior of the APC-binding protein EB1 on the distal ends of microtubules. Curr. Biol. 10, 865–868.

Moissoglu, K., Stueland, M., Gasparski, A.N., Wang, T., Jenkins, L.M., Hastings, M.L., and Mili, S. (2020). RNA localization and co-translational interactions control RAB 13 GTP ase function and cell migration. EMBO J. 39.

Monroy, B.Y., Sawyer, D.L., Ackermann, B.E., Borden, M.M., Tan, T.C., and Ori-Mckenney, K.M. (2018). Competition between microtubule-associated proteins directs motor transport. Nat. Commun. 9, 1–12.

Monroy, B.Y., Tan, T.C., Oclaman, J.M., Han, J.S., Simó, S., Niwa, S., Nowakowski, D.W., McKenney, R.J., and Ori-McKenney, K.M. (2020). A Combinatorial MAP Code Dictates Polarized Microtubule Transport. Dev. Cell 53, 60-72.e4.

Park, H.Y., Lim, H., Yoon, Y.J., Follenzi, A., Nwokafor, C., Lopez-Jones, M., Meng, X., and Singer, R.H. (2014). Visualization of dynamics of single endogenous mRNA labeled in live mouse. Science 343, 422–424.

Preitner, N., Quan, J., Nowakowski, D.W., Hancock, M.L., Shi, J., Tcherkezian, J., Young-Pearse, T.L., and Flanagan, J.G. (2014). APC Is an RNA-Binding Protein, and Its Interactome Provides a Link to Neural Development and Microtubule Assembly. Cell 158, 368–382.

Reilein, A., and Nelson, W.J. (2005). APC is a component of an organizing template for cortical microtubule networks. Nat. Cell Biol. 7, 463–473.

Rodrigues, E.C., Grawenhoff, J., Baumann, S.J., Lorenzon, N., and Maurer, S.P. (2021). Mammalian Neuronal mRNA Transport Complexes: The Few Knowns and the Many Unknowns. Front. Integr. Neurosci. 15, 1–8.

Ruane, P.T., Gumy, L.F., Bola, B., Anderson, B., Wozniak, M.J., Hoogenraad, C.C., and Allan, V.J. (2016). Tumour Suppressor Adenomatous Polyposis Coli (APC) localisation is regulated by both Kinesin-1 and Kinesin-2. Sci. Rep. 6, 27456.

Scarborough, E.A., Uchida, K., Vogel, M., Erlitzki, N., Iyer, M., Phyo, S.A., Bogush, A., Kehat, I., and Prosser, B.L. (2021). Microtubules orchestrate local translation to enable cardiac growth. Nat. Commun. 12.

Schindelin, J., Arganda-Carreras, I., Frise, E., Kaynig, V., Longair, M., Pietzsch, T., Preibisch, S., Rueden, C., Saalfeld, S., Schmid, B., et al. (2012). Fiji: an open-source platform for biological-image analysis. Nat. Methods 9, 676–682.

Scholz, J., Besir, H., Strasser, C., and Suppmann, S. (2013). A new method to customize protein expression vectors for fast, efficient and background free parallel cloning. BMC Biotechnol. 13, 12.

Sharangdhar, T., Sugimoto, Y., Heraud-Farlow, J., Fernández-Moya, S.M., Ehses, J., Ruiz de Los Mozos, I., Ule, J., and Kiebler, M.A. (2017). A retained intron in the 3’-UTR of Calm3 mRNA mediates its Staufen2- and activity-dependent localization to neuronal dendrites. EMBO Rep. 18, 1762–1774.

Song, M.S., Moon, H.C., Jeon, J.H., and Park, H.Y. (2018). Neuronal messenger ribonucleoprotein transport follows an aging Lévy walk. Nat. Commun. 9, 1–8.

Su, L.K., Burrell, M., Hill, D.E., Gyuris, J., Brent, R., Wiltshire, R., Trent, J., Vogelstein, B., and Kinzler, K.W. (1995). APC binds to the novel protein EB1. Cancer Res. 55, 2972–2977.

Tinevez, J.Y., Perry, N., Schindelin, J., Hoopes, G.M., Reynolds, G.D., Laplantine, E., Bednarek, S.Y., Shorte, S.L., and Eliceiri, K.W. (2017). TrackMate: An open and extensible platform for single-particle tracking. Methods 115, 80–90.

Tushev, G., Glock, C., Heumüller, M., Biever, A., Jovanovic, M., and Schuman, E.M. (2018). Alternative 3’ UTRs Modify the Localization, Regulatory Potential, Stability, and Plasticity of mRNAs in Neuronal Compartments. Neuron 98, 495-511.e6.

van Haren, J., Boudeau, J., Schmidt, S., Basu, S., Liu, Z., Lammers, D., Demmers, J., Benhari, J., Grosveld, F., Debant, A., et al. (2014). Dynamic Microtubules Catalyze Formation of Navigator-TRIO Complexes to Regulate Neurite Extension. Curr. Biol. 24, 1778–1785.

Viswanathan, G.M., Buldyrev, S. V, Havlin, S., da Luz, M.G.E., Raposo, E.P., and Stanley, H.E. (1999). Optimizing the success of random searches. Nature 401, 911–914.

Wang, G., Ang, C., Fan, J., Wang, A., Moffitt, J.R., and Zhuang, X. (2020). Spatial organization of the transcriptome in individual neurons. BioRxiv 1–45.

Wong, H.H.-W., Lin, J.Q., Ströhl, F., Roque, C.G., Cioni, J.-M., Cagnetta, R., Turner-Bridger, B., Laine, R.F., Harris, W.A., Kaminski, C.F., et al. (2017). RNA Docking and Local Translation Regulate Site-Specific Axon Remodeling In Vivo. Neuron 95, 852-868.e8.

Yoon, Y.J., Wu, B., Buxbaum, A.R., Das, S., Tsai, A., English, B.P., Grimm, J.B., Lavis, L.D., and Singer, R.H. (2016). Glutamate-induced RNA localization and translation in neurons. Proc. Natl. Acad. Sci. U. S. A. 113, E6877–E6886.

Zhao, J., Fok, A.H.K., Fan, R., Kwan, P.-Y., Chan, H.-L., Lo, L.H.-Y., Chan, Y.-S., Yung, W.-H., Huang, J., Lai, C.S.W., et al. (2020). Specific depletion of the motor protein KIF5B leads to deficits in dendritic transport, synaptic plasticity and memory. Elife 9, 1–34.

Zumbrunn, J., Kinoshita, K., Hyman, A.A., and Näthke, I.S. (2001). Binding of the adenomatous polyposis coli protein to microtubules increases microtubule stability and is regulated by GSK3β phosphorylation. Curr. Biol. 11, 44–49.

