## Supplemental data for "APC couples neuronal mRNAs to multiple kinesins, EB1 and shrinking microtubule ends for bidirectional mRNA motility"

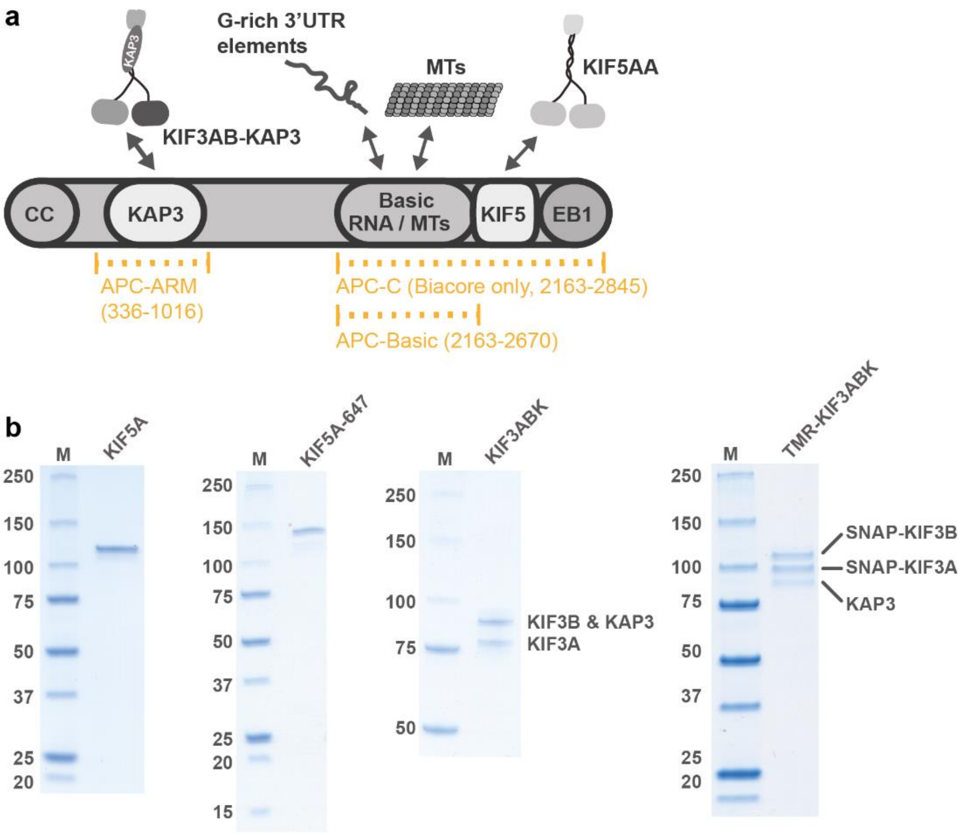


c

| Name | Sequence | Accession Number | Label | Length (nt) | % G | longest G-stretch | dG (kcal/mol) |
|---|---|---|---|---|---|---|---|
| β2Btub$_{wt}$ | GUUCCCCAGGCCAAGCAGGUUAGGGAAAGCUGAGAUGAAAGGAGGGGGUGGGGGGGCUUAAUCUGUGAAAAUACCUUGGCAGUUGAA | NM_023716.2 | 5' Alexa-647 | 87 | 38 | 6 | -23.1 |
| β2Btub$_{wtmin}$ | GAAAGGAGGGGGUGGGGGGGCUUAA | NM_023716.2 | - | 25 | 60 | 6 | -0.6 |
| β2Btub$_{mutmin}$ | AGUACGUCCUUGGCUAGAGAUAGAC | N/A | - | 25 | 28 | 2 | -1.3 |
| β-actin | UGGGAGGGGGUUGAGGUGUUGAGGC | NM_007393.5 | - | 25 | 60 | 5 | 1.3 |
| Camk2α | CGGATGGGTTGGGAAGTGGGTGGGA | NM_001286809.1 | - | 25 | 60 | 3 | 2.1 |
| Map1b | GCGGGAGGGGAGGGAGAGAGGCGGG | NM_008634.2 | - | 25 | 72 | 4 | 0.4 |

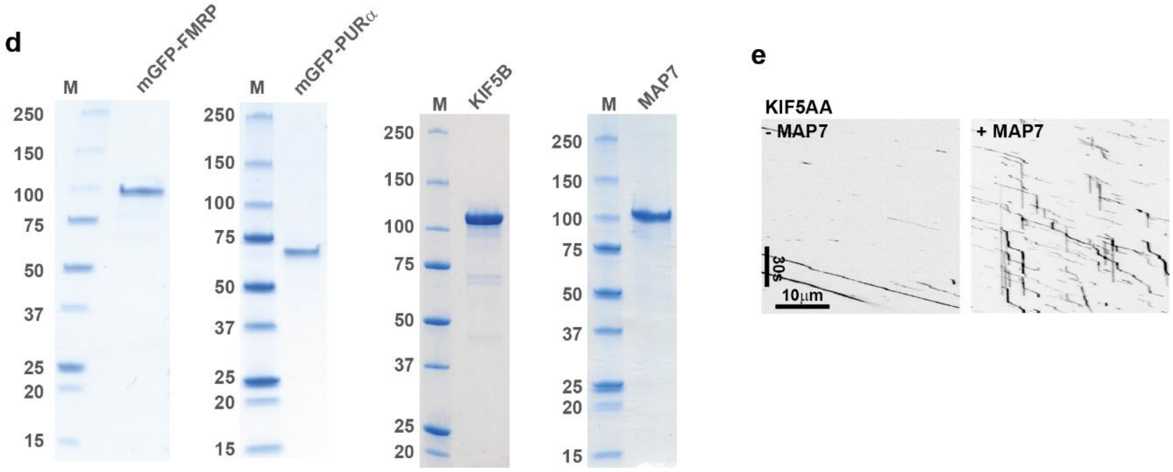


**Supplementary Fig. 1. (a)** Cartoon depicting known APC interaction sites relevant for this work. CC: Coiled coil domain for APC dimerisation, KAP3: Cargo-binding adaptor of kinesin-2, Basic domain: Interacts with the microtubule lattice and RNAs, KIF5: Kinesin-1-binding site, EB1: EB1-binding site. **(b)** Coomassie-stained SDS-gels of purified, recombinant KIF5A, Alexa647-labelled KIF5A, unlabelled heterotrimeric kinesin-2 (note: KIF3B and KAP3 migrate together, which has been demonstrated previously by SDS-PAGE and Western blotting (Baumann et al., 2020)) **(c)** Table of mRNA fragments used in this work. **(d)** Coomassie-stained SDS-gels of purified, recombinant mGFP-FMRP, mGFP-PURα, KIF5B and MAP7. **(e)** Kymographs of 75 pM KIF5AA-A647 in the absence and presence of 10000 pM MAP7.

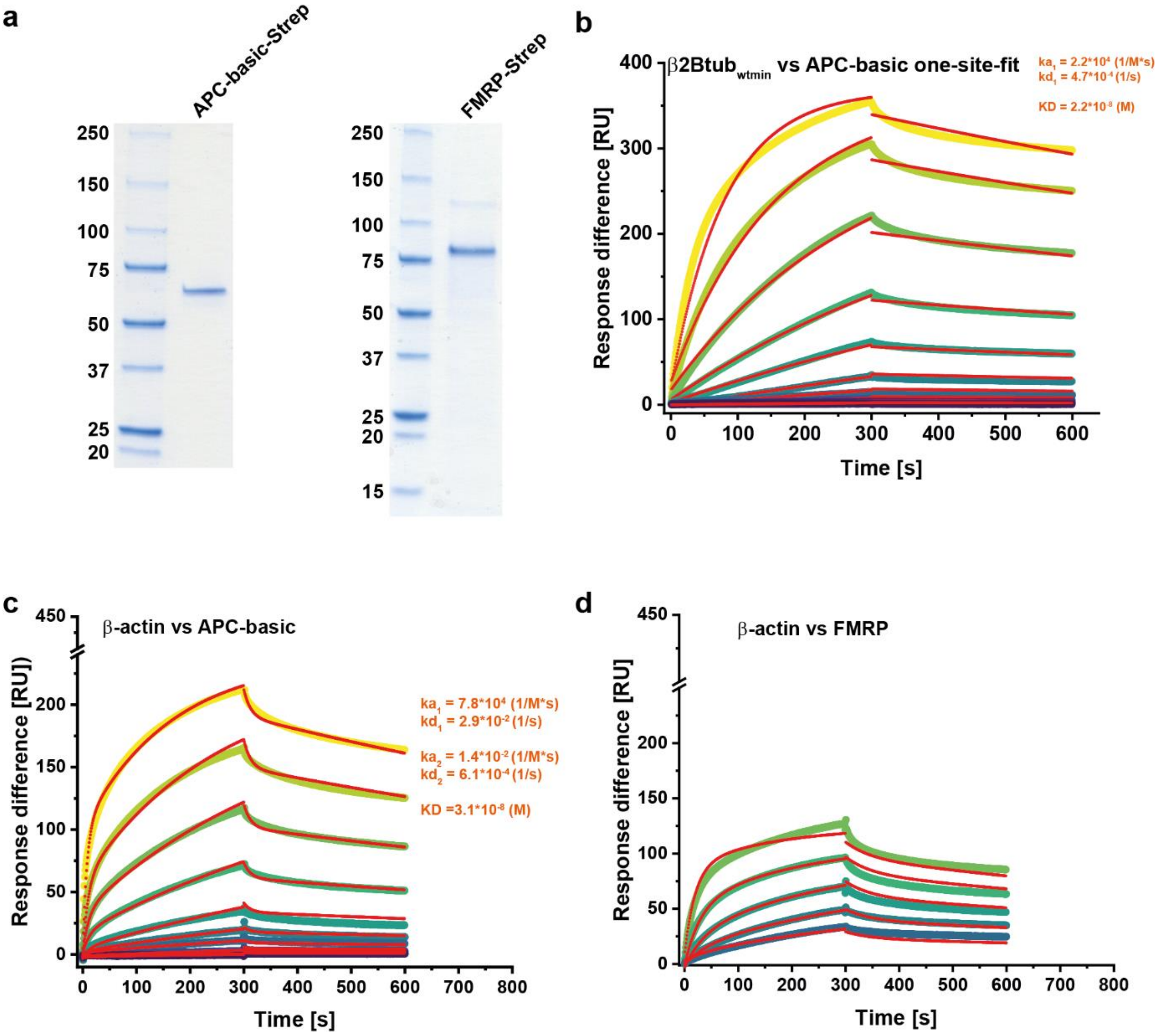


**Supplementary Fig. 2. (a)** Coomassie-stained SDS-gels of C-terminally Strep-tagged APC-basic domain and full-length FMRP. **(b)** Same graph as in Fig.2a. with a one-site fit model. **(c & d)** Graphs generated from Biacore experiments showing the binding kinetics (association and dissociation rate) of different concentrations of β2Btub$_{wtmin}$ RNA and APC-basic domain or FMRP, respectively. Experimentally determined curves were fitted with a two-site fit model.

**a**

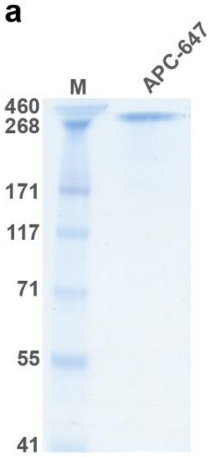


**Supplementary Fig. 3. (a)** Coomassie-stained SDS-gels of APC-C-SNAP labelled with Alexa647.

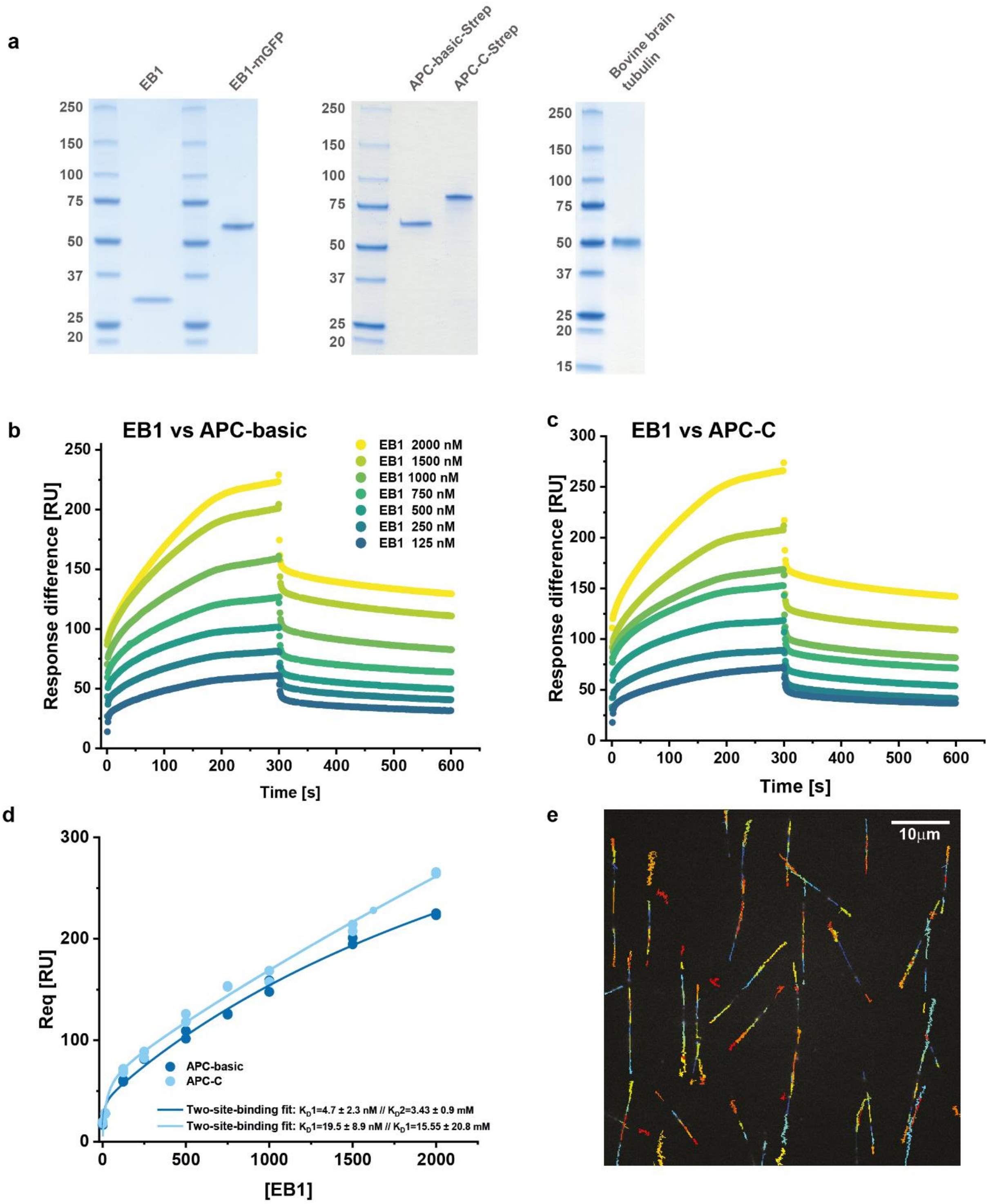


**Supplementary Fig. 4. (a)** Coomassie-stained SDS-gels of EB1, EB1-mGFP, APC-basic-Strep, APC-C-Strep and bovine tubulin. **(b & c)** Graphs generated from Biacore experiments showing the binding kinetics (association and dissociation rate) of different concentrations of EB1 and APC-basic as well as APC-C, respectively. **(d)** Binding curves showing the interaction of EB1 with immobilised APC-basic and APC-C fragments. A two-site binding model was used to fit the Bmax-derived Req-values. **(e)** Capture of an EB1-mGFP tracking experiment with overlain tracks. Shown is the whole field of view 512 x 512 pixels. Tracks identified by Trackmate (ImageJ) are colour-coded according to their chronological appearance (blue = early, red = late). Duration of the experiment is 450 seconds. The overlap of tracks results from repeated growth and shrinking cycles. Only growing MT ends are detected.
